# Motor dysfunction in *Drosophila melanogaster* as a biomarker for developmental neurotoxicity

**DOI:** 10.1101/2021.07.09.451676

**Authors:** Ana Cabrita, Alexandra M. Medeiros, Telmo Pereira, António Sebastião Rodrigues, Michel Kranendonk, César S. Mendes

## Abstract

Humans interact with numerous chemical compounds with direct health implications, with several able to induce developmental neurotoxicity (DNT), which bear developmental, behavioral, and cognitive consequences from a young age. Current guidelines for DNT testing are notably costly, time consuming, and unsuitable for testing large numbers of chemicals. Therefore, there is a need for adequate alternatives to conventional animal testing for neurotoxicity and DNT. Here we show that detailed kinematic analysis can provide a strong indicator for DNT, using known (chlorpyrifos, CPS) or putative (β-N-methylamino-L-alanine, BMAA) neurotoxic compounds. We exposed *Drosophila melanogaster* to these compounds during development and evaluated for common general toxicity — notably developmental survival and pupal positioning, together with the FlyWalker system, a detailed adult kinematics evaluation method.

At concentrations that do not induce general toxicity, the solvent DMSO had a significant effect on kinematic parameters. Nonetheless, CPS not only induced developmental lethality but also significantly impaired coordination in comparison to DMSO, altering 16 motor parameters, validating the usefulness of our kinematic approach.

Interestingly, BMAA, although not lethal during development, induced a dose-dependent motor decay, targeting most parameters in young adult animals, phenotypically resembling normally aged, non-exposed flies. This effect was subsequently attenuated during ageing, indicating an adaptive response. Furthermore, BMAA induced an abnormal terminal differentiation of leg motor neurons, without inducing degeneration, underpinning the observed altered mobility phenotype. Overall, our results support our kinematic approach as a novel, highly sensitive and reliable tool to assess potential DNT of chemical compounds.

## Introduction

Neurotoxicity and developmental neurotoxicity (DNT) are important adverse health effects of environmental contaminants, occupational chemicals, and natural toxins. Moreover, neurotoxicity is one of the most frequent occurring therapeutic drug side effect (Bal-Price et al., 2015b). Exposure to neurotoxicants during development has been recognized to be of particular importance as the developing human brain is inherently more susceptible to damaging agents than is the brain of an adult (Bondy and Campbell, 2005; Rodier, 1995). DNT has been implicated in the etiology of various neuropsychiatric and neurological disorders, including autism spectrum disorder, attention deficit hyperactive disorder, schizophrenia, Parkinson’s, and Alzheimer’s disease (Grandjean and Landrigan, 2014).

Many cellular and molecular processes are known to be crucial for a proper development and function of the central (CNS) and peripheral nervous systems (PNS). However, there are relatively few examples of well-documented pathways how chemicals may interfere in these processes (Bal-Price and Meek, 2017). The paucity of toxicological data for tens of thousands of chemicals in commercial use and thousands of new chemicals produced each year, has called the attention during the last decade, for development of sensitive and rapid assays to screen for their neurotoxic propensity, particular for DNT (Smirnova et al., 2014).

The main reason for the lack of data lies in the current guidelines (OECD TG 426 and US EPA 712-C-98-239; OECD, 2007; US EPA, 1998) largely based on in vivo experiments, which are costly, time consuming, and unsuitable for testing large numbers of chemicals (Bal-Price et al., 2012). Therefore, there is a need for adequate alternatives to conventional animal testing for neurotoxicity and DNT (Bal-Price *et al*., 2012). Hence efforts are being directed toward the development of alternative models, utilizing either mammalian cells in culture or non-mammalian model systems, with the inclusion of new testing strategies to facilitate transition to more mechanistically based approaches (Xie et al., 2020). Such approaches can unveil mechanistic cues that can assist in optimizing sensitive endpoints for pathway-specific screening and the determination of predictive key events, which can be used as biomarkers for specific neurotoxic outcomes (Bal-Price and Meek, 2017). Models such as zebra fish, *C. elegans* and *Drosophila melanogaster* are increasingly recognized for their suitability to test a larger number of candidate compounds, in addition to acquire “mode of action” type of information regarding neurotoxicity, and neurological disorders and diseases (Peterson et al., 2008). The fruit fly *Drosophila melanogaster* has got particular attention in the modern regimen of neurotoxicological testing due to similarities of their neurological and developmental pathways with those of vertebrates (Rand, 2010), emphasizing its unique attributes for assaying neurodevelopment and behavior (Affleck and Walker, 2019; Rand et al., 2019). Recent investigations have propagated several powerful assay methods with *Drosophila* in developmental and behavioral toxicology (Rand, 2010). However, most studies rely on parameters that poorly reflect neuronal defects or decay. In these studies, several parameters are tested in an attempt to find parameters indicative of more general toxicity such as mortality, female-male ratio, deoxyribonucleic acid (DNA) damage (Cox et al., 2016), climbing assay, alteration in acetylcholinesterase activity and other enzymes (Rand et al., 2014). Despite their usefulness, climbing assays only provide low resolution in distinguishing different experimental conditions, i.e., just a one-dimensional perspective of a complex and multidimensional phenomenon such as multi-jointed walking, highly dependent on a properly developed and fine-tuned nervous system. Coordinated walking in vertebrates and multi-legged invertebrates such as *D. melanogaster* requires a complex neural network coupled to sensory feedback (Dickinson et al., 2000). Perturbation of such neural networks in humans has been indicated as the source of several neurological disorders, in which exposure to neurotoxic chemicals was implicated (Grandjean and Landrigan, 2014).

The combination of increasingly sophisticated optical systems with computer algorithms have allowed to track movement of multi-segmented body parts with high spatio-temporal resolution and extract a multitude of quantifiable data that accurately describe walking performance (Berman et al., 2014; Günel et al., 2019; Kain et al., 2013; Mathis et al., 2018; Mendes et al., 2013; Pereira et al., 2019; Uhlmann et al., 2017; Wu et al., 2019). Such tools have identified the effects on the motor system of internal and external manipulations such as increased body load or lack of a particular neurotransmitter, amongst others (Enriquez et al., 2015; Howard et al., 2019; Mendes et al., 2014; Schretter et al., 2018).

In the current report we describe the use of *Drosophila melanogaster* as a model to study the DNT of a known neurotoxic agent standardly applied in testing, namely the pesticide chlorpyrifos (CPS) (Aschner et al., 2017), in addition to testing the putative neurotoxic agent β-N-methylamino-L-alanine (BMAA) (Weiss and Choi, 1988). For this we combined classical metrics such as animal survival with a sensitive kinematic assay that accurately and quantitatively reports the status of the motor system (Mendes *et al*., 2013), using it as indicator of DNT.

## Results

### Effect of DMSO and CPS on developmental survival

CPS, a commonly used organophosphate pesticide and a known neurotoxin applied as a standard in neurotoxicity screenings, was used to study its effect on the development of *Drosophila melanogaster,* by transferring fertilized eggs into food containing defined concentrations of this compound (Fig. 1A).

**Fig. 1.**
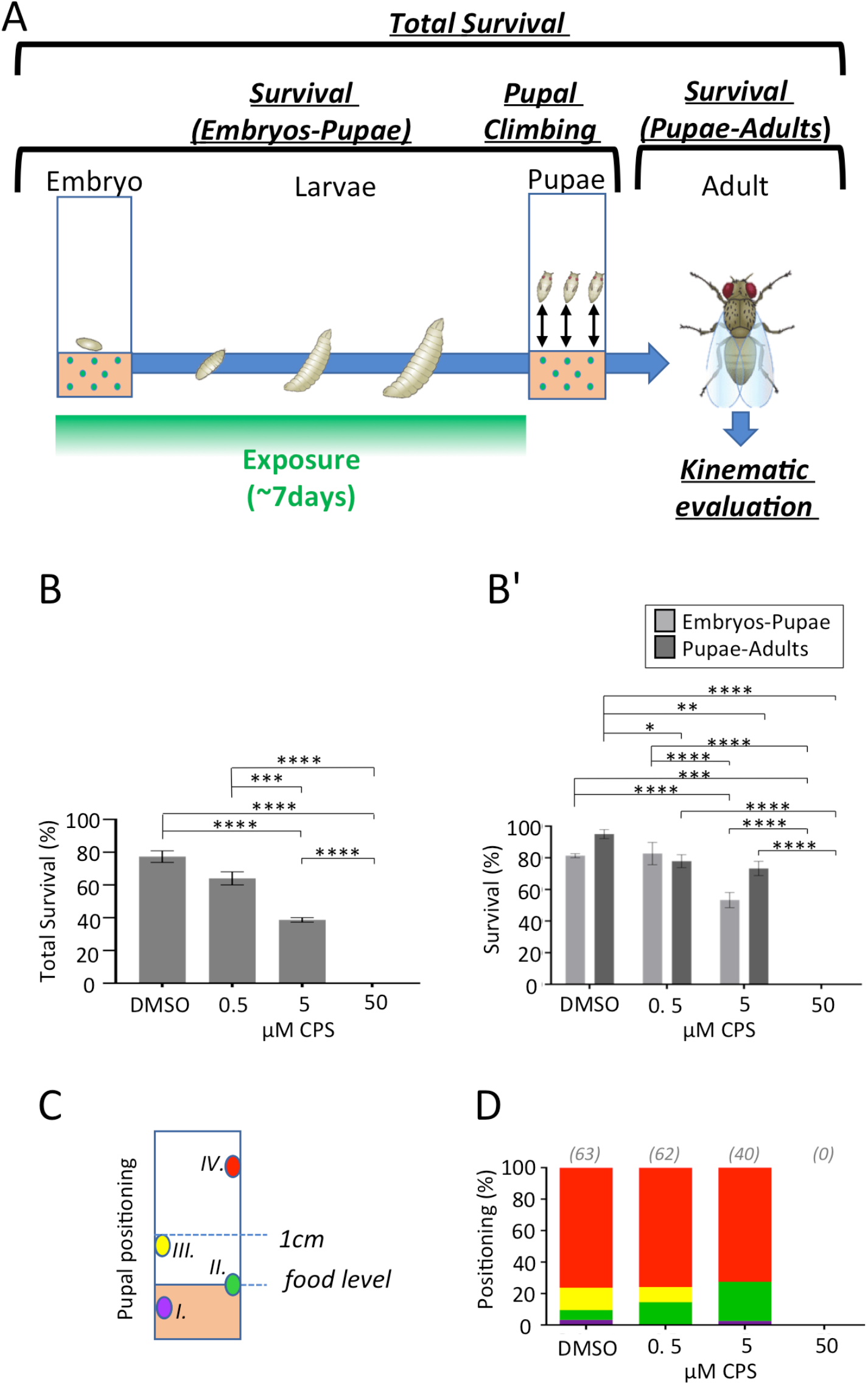
Measurements of pupal and adult survival of *Drosophila melanogaster* developing in the presence of the neurotoxic agent CPS. (A) Schematic of the procedure and metrics used in this study. Embryos are placed in tainted or untainted food (control). Several parameters were quantified in this study (in italic), see main text for details. (B) Survival of adult animals, which were exposed during development to increasing concentrations of CPS (0; 0.5; 5 or; 50 µM; n = 75 for each condition) compared to solvent (DMSO). Bar graphs represent the average percentage of animals surviving each developmental stage ± SEM. Statistical analysis with two-way ANOVA followed by Tukey’s post hoc test, *P < 0.05; **P < 0.01; ***P < 0.001. (B’) Partial survival scores. Light grey represent embryo to pupae survival, and dark grey represent pupae to adult survival. (C-D) Pupal distance. (C) Climbing scoring scheme: Pupal positions were divided into four groups: I. (purple) inside the food; II. (green) slightly above the food; III. (yellow) up to 1cm from the food; and IV. (red) occupying the area above 1 cm. (D) Pupal distance represented as a percentage stacked bar graph for increasing concentrations of CPS compared to solvent (DMSO). Number of animals tested (n) is indicated in the figure.

*Drosophila*’s development from an egg to adult takes approximately 11 days at 25°C, comprising a stationary embryonic stage, lasting 22 hrs. (Ashburner, 1989), followed by three larval stages, in which activity consists of mostly food consumption with a duration of four days. Subsequently, third instar larvae crawl away from their food source and start a molting process outside the food-source, undergoing a metamorphosis process lasting approximately 3 days (Denlinger and Zdarek, 1994; Markow et al., 2009; Powsner, 1935). The development of any multicellular organism depends on the rigorous execution of a specific developmental and genetic program. As this developmental process is highly sensitive to chemical insults, we measured initially three developmental features: i) survival from egg to adult; ii) egg to pupae; and iii) pupae to adult (Fig. 1A), an approach described previously (Khatun et al., 2018; Nazir et al., 2001; Zhou et al., 2010b). Due to poor aqueous solubility, CPS was diluted in aqueous DMSO, which although is used widely as solvent in experimental biology and a preferred cryoprotectant, can *per se* induce developmental toxicity (Gurtovenko and Anwar, 2007; Nazir et al., 2003; Uysal et al., 2015) and neurotoxicity (Awan et al., 2020; Bakar et al., 2012). Control experiment were performed, testing several food concentrations of DMSO at pupal and adult stages, to determine a concentration that would allow to predilute CPS, without interference of this solvent (Fig. S1; see Methods). A concentration of 140 mM DMSO in fly food induced a reduced number of eclosed animals (hatching from the pupal case), consequence of a reduced pupal to adult transition (Fig. S1B). However, lower concentrations of DMSO displayed little or no effect on survival, and a 70 mM aqueous solution of DMSO was subsequently used as solvent for CPS (Fig. S1B). Exposure of developing *Drosophila* to increasing concentrations of CPS during development showed a dose-dependent lethality with concentrations as low as 5 µM showing an approximately 50% reduction in the number of animals reaching adulthood (Fig. 1B). Additional drop in survival (approximately 22%) was found during the pupa to adult transition (Fig. 1B’), indicating the detrimental effect of CPS even after animals ceased to be exposed to the tainted food. A 50 µM concentration of CPS induced complete lethality with no animals reaching the pupal stage (Fig.1B’).

In *Drosophila*, the feeding larval stage is followed by a wondering phase where larvae exit food to find an appropriate pupation site (Denlinger and Zdarek, 1994). The pupation travelling distance can be influenced by genetic traits and external factors such as moisture and environmental cues (Beltramí et al., 2010; Johnson and Carder, 2012; Narasimha et al., 2015; Sokolowski and Bauer, 1989). In addition, the distance to the pupation site has been used as a marker for fitness of the neuromuscular system to sense the surrounding environment and drive the crawling larva away from the food source to a safe location for the metamorphosis process to occur (Johnson and Carder, 2012; Joshi and Mueller, 1993), notably upon exposure of the developing larva to toxic compounds (Khatun *et al*., 2018; Sood et al., 2019). Accordingly, we measured the pupation distance by recording the position of pupas relative to the food level in four categories: I. within the food; II. slightly above the food line; III. up to 1 cm from the food; and IV. above 1 cm mark (Fig. 1C). The pupal positioning of developing animals exposed to increasing concentrations of CPS was subsequently monitored, according to these positioning categories (Fig. 1D and Table S1). Increasing concentrations of CPS did not significantly change the percentage of pupae located beyond the 1 cm mark (76 ± 10 % for control condition vs 76 ± 7 % and 73 ± 1 %, for 0.5 and 5 µM, respectively; Fig. 1D and Table S1). Moreover, there was an increase, albeit non-significant, in the number of pupae present at the food interface, simultaneously matched by a decrease of pupa above the food line but below the one cm mark (Fig. 1D and Table S1). Although statistical analysis did not show statistical significance between these groups, these data suggest that CPS can affect the ability of wondering larva to emerge from the food, in order to pupate. Overall, these results show that the known neurotoxic agent –CPS-can affect the ability of *Drosophila* to survive development.

### Effect of DMSO and CPS on adult kinematics

Surviving adult animals that developed in the presence of CPS did not show clear anatomical defects (data not shown). Subsequently, we tested if there were any detrimental effects of this known neurotoxin on coordination in their walking conduct (Fig. 1A). Walking behavior of multi-segmented organisms such as *Drosophila* display a highly reproducible and stereotyped pattern aimed to move the animal in an energy-efficient and stable fashion (Mendes *et al*., 2013; Ramdya et al., 2017; Strauss and Heisenberg, 1990; Szczecinski et al., 2018; Wosnitza et al., 2013). However, disruption of the motor neuronal circuit has direct consequences on gait properties of walking animals. For example, inactivation of serotonergic neurons in the ventral nerve cord of *Drosophila* (the analog of the mammalian spinal cord) has a significant effect on gait properties (Howard *et al*., 2019). We thus considered that exposure to neurotoxins may not lead to lethality during development but nevertheless could affect the proper development and function of the motor nervous system. To test this hypothesis, we analyzed the kinematic behavior of untethered adult flies, which during development were exposed to food containing non-lethal levels of CPS (5 µM) using the FlyWalker system. This approach allows the extraction of a large set of kinematic parameters with a high spatiotemporal resolution (Mendes *et al*., 2013) (Fig. 2 and S2).

**Fig. 2.**
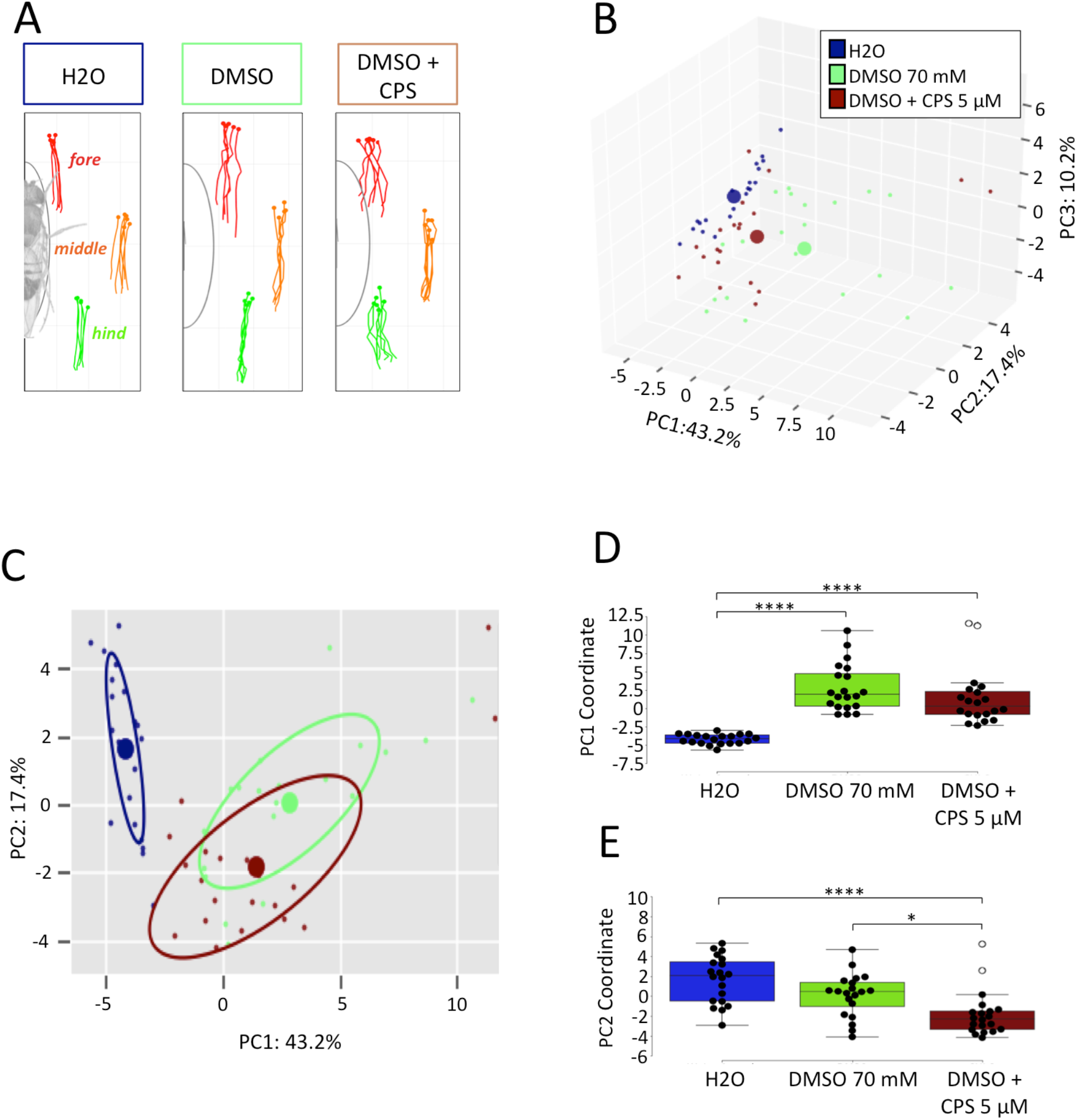
Kinematic parameters of adult animals exposed during development to water, 70 mM DMSO (solvent) and DMSO plus 5 µM CPS. One-week old animals were exposed during development to water, 70 mM DMSO (solvent) and DMSO plus 5 µM CPS and kinematics features were recorded and analyzed using the FlyWalker system (n=20 for each condition). (A) Representative stance traces, which marks the tarsal contacts relative to the body axis during stance phases. See (Mendes *et al*., 2013) for details. (B-D) PCA of all kinematic parameters. (B) Tridimensional representation of three-component PCA analysis. Each individual small dot represents one video while larger dots represents the average point (n=20 for each condition). Contribution of each component is indicated in each axis; (C) 2D representation with circles representing 50% of collected data; (D-E) Comparison of each PC coordinate. Boxplots represent the median as the middle line, with the lower and upper edges of the boxes representing the 25% and 75% quartiles, respectively; the whiskers represent the range of the full data set, excluding outliers (open circles). *P < 0.05; **P < 0.01; ***P < 0.001. (D) PC1 coordinates. (E) PC2 coordinates. See Methods for details.

We compared the kinematic performance of adult animals developing in the presence of water, 70 mM of the solvent DMSO, and the solvent plus 5 µM CPS (Fig. 2, S2 and S3). When compared to water, animals developing in the presence of the aqueous DMSO alone or in combination with 5 µM CPS displayed stance phases with inconsistent leg positioning relative to the body axis, including during touch-down and swing-onset (Fig. 2A). Consistently, by analyzing the kinematic features of our experimental conditions, we found that solvent alone or in the presence of CPS did affect a significant number of parameters relative to water (Fig. S2B). Moreover, comparing the performance of 70 mM DMSO to DMSO plus 5 µM CPS, we found that CPS induced kinematic alterations on 16 kinematic parameters (Fig. S2B), most notably on step frequency, step length and stance straightness, which measures the consistency of the stance phase (or power stroke), indicating that CPS promotes an aggravated uncoordinated walking behavior compared to its solvent. We also subjected all the kinematic parameters to a three-order Principal Component Analysis (PCA), generating a three-dimensional representation (Fig. 2 B-E). PC1, with a 43,2% variance in the data set, allowed the discrimination of exposure between water vs. DMSO and vs. DMSO plus 5 µM CPS (Fig. 2B-D). However PC2, with a 17.4% variation variance, allowed the differentiation of animals exposed to CPS vs. 70 mM DMSO alone (Fig. 2C,E), further indicating a kinematic effect induced by the exposure to CPS during development.

**Fig. 3.**
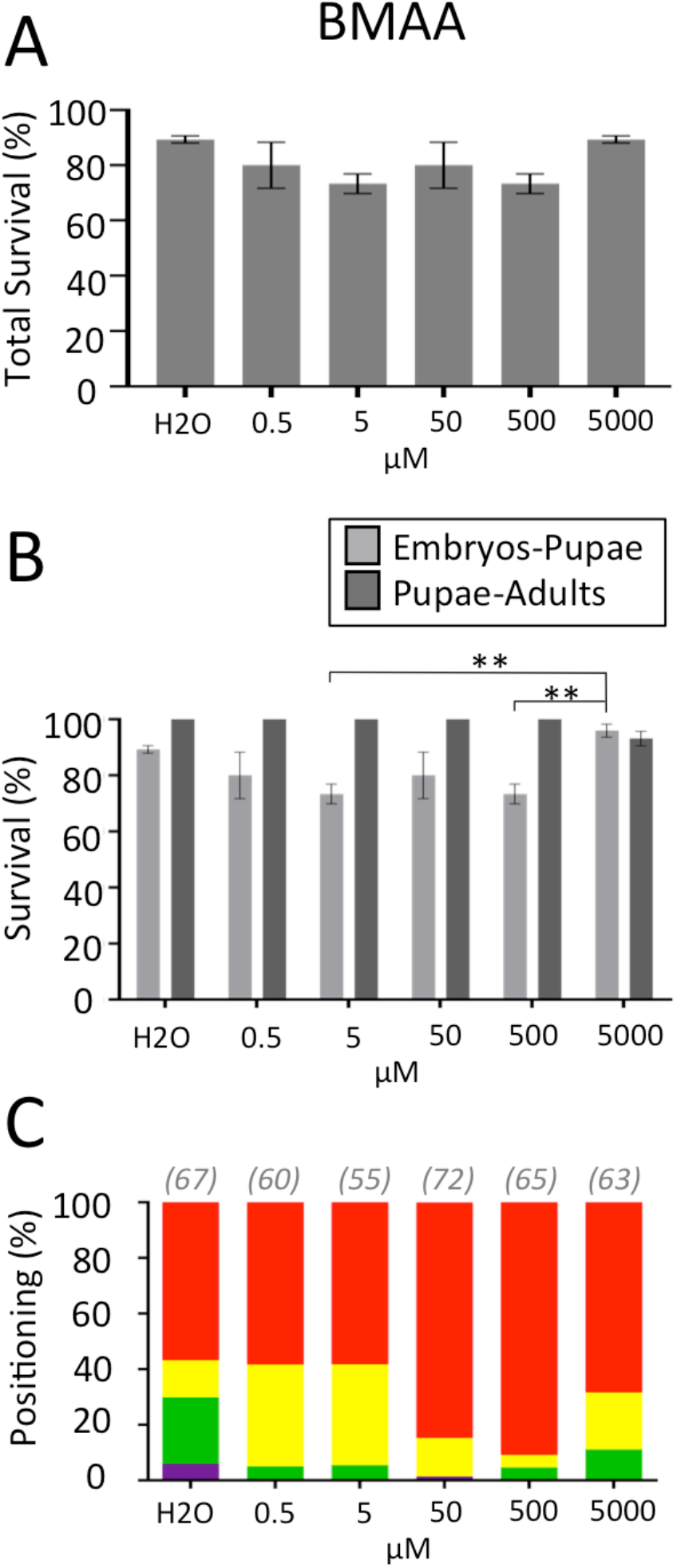
Pupal and adult survival of *Drosophila melanogaster* developing in the presence of the BMAA. (A) survival of adult animals developing in increasing concentrations of BMAA (0; 0.5; 5 or 500 µM; n =75 for each condition) compared to solvent (water). (B) Partial survival scores. Light grey represent embryo to pupae survival, and dark grey represent pupae to adult survival. Bar graphs represent the average percentage of animals surviving each developmental stage ± SEM. Statistical analysis with two-way ANOVA followed by Tukey’s post hoc test. (C) Pupal distance. Pupal positions were divided into four groups: I. (purple) inside the food; II. (green) slightly above the food; III. (yellow) up to 1cm from the food; and IV. (red) occupying the area above 1 cm. Number of animals tested (n) is indicated in the figure. Results represented as a percentage stacked bar graph for increasing concentrations BMAA compared to solvent (water).

Altogether, our results supports that animals surviving exposure to specific levels of neurotoxic agents during development may still display motor dysfunction, highlighting the presence of neuronal defects. Our data indicates that DMSO can mask DNT effects in *Drosophila*, which can hamper an adequate assessment of test compounds. This is an important issue for *in vitro* toxicological studies in general regarding solvents (Coecke et al., 2016). Nevertheless, our data analysis highlighted the ability of our Flywalker approach in a quantitative manner to identify CPS, capable to induce kinematic dysfunction and thus developmental neurotoxicity.

### Effect of BMAA on developmental survival

Subsequently, we evaluated the effect of BMAA on the development of *Drosophila*. The non-proteinogenic amino acid BMAA has been indicated to induce neurotoxic effects and has been implicated in the etiology of amyotrophic lateral sclerosis (ALS) (Banack et al., 2015; Jonasson et al., 2010; Lobner et al., 2007; Muñoz-Sáez et al., 2015; Murch et al., 2004; Pablo et al., 2009; Proctor et al., 2019; Roy-Lachapelle et al., 2017; Weiss et al., 1989). Former studies of BMAA in *Drosophila* showed that BMAA severely reduced life span, climbing capabilities and memory (Zhou et al., 2009; Zhou et al., 2010a). Interestingly, our data shows that exposure to increasing food concentrations of BMAA did not affect the survival of developing *Drosophila*, with similar number of animals reaching adulthood and pupal stages compared to control animals (solvent, i.e. water) (Fig. 3A, B). Still, BMAA induced a shift in pupal positioning from positions below or at the food level, a strong indication for larval neuromuscular system impairment, to positions located further away from the food source (Fig. 3C, Table S2), suggesting a potential repellent or escaping effect of BMAA on wondering third instar larvae.

### Effect of BMAA on adult kinematics

Although BMAA did not affect developmental viability, we tested if developing *Drosophila* exposed to increasing concentrations of this amino acid displayed motor dysfunction, using the FlyWalker system. We found that 1-week old adult *Drosophila* displayed a dose-depended motor dysfunction when exposed to increasing concentration of BMAA during development (Fig. 4, S4). Stance phases display an increasingly inconsistent leg positioning relative to the body axis, including during touch-down and swing-onset (Fig. 4A). Remarkably, all parameter classes were altered, although step length was not affected at any of the tested concentrations, suggesting that this gait feature is insensitive to developmental dysfunction induced by BMAA (Fig. 4B). The PCA analysis indicated a 37.1% variance of data in PC1, and that increasing concentrations of BMAA corresponded to a stronger effect (i.e. dose-response), relative to the control condition (Fig. 4C-E). However, PC2 representing 17.6% variance did not allow any discrimination between control and experimental groups (Fig. 4F), indicating the capture of the most relevant variation in motor function by PC1. Similarly, individual analysis of gait parameters confirmed the aforementioned trend with several parameters showing increasing divergence from control conditions with increased concentrations of BMAA (Fig. S4).

**Fig. 4.**
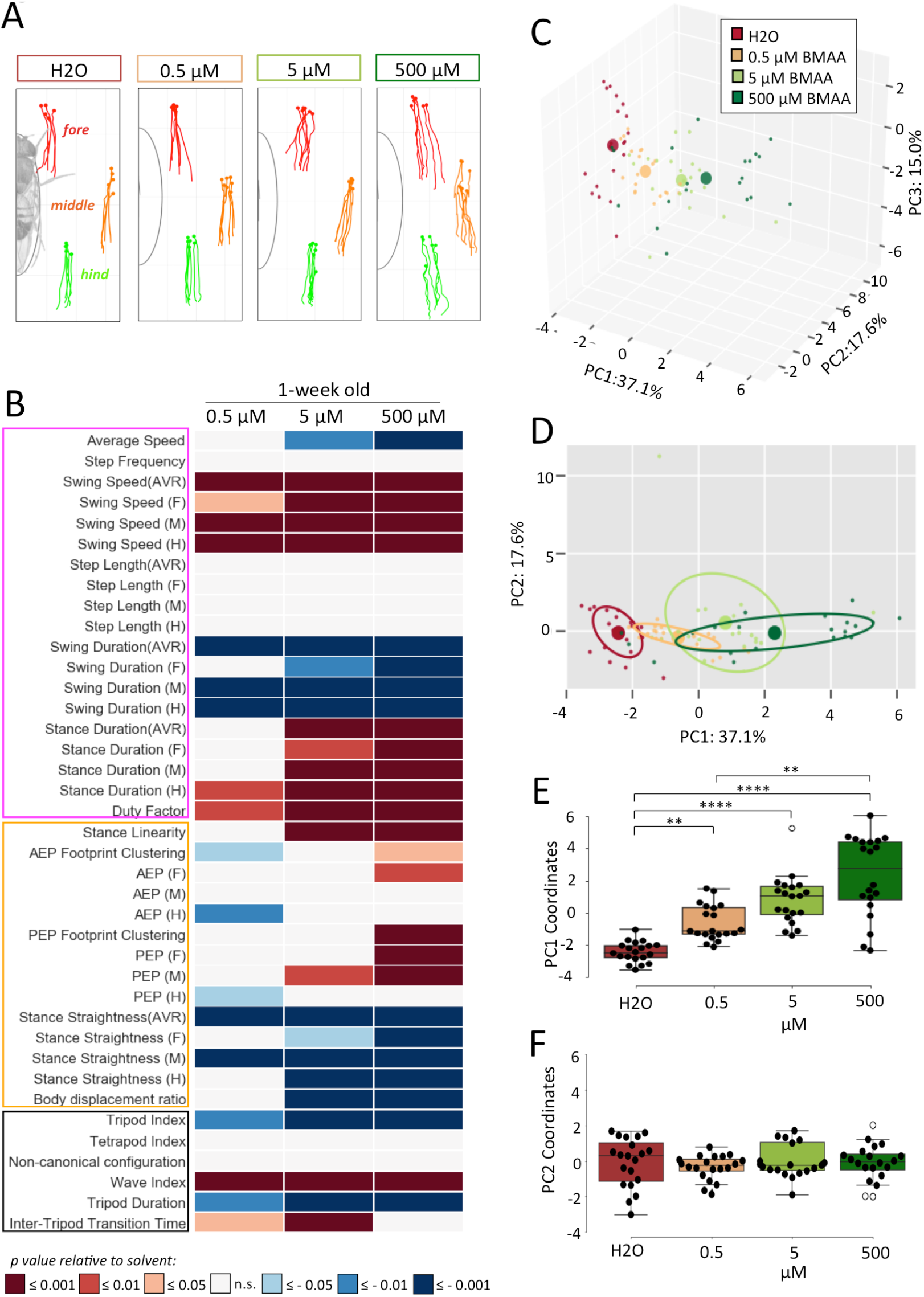
Kinematic parameters of one-week adult animals exposed to increasing concentrations of BMAA during development. Exposure levels were 0; 0.5; 5 or 500 µM of BMAA. Kinematic features were recorded and analyzed using the FlyWalker system (n=20 for each condition). (A) Representative stance traces, which marks the tarsal contacts relative to the body axis during stance phases. See (Mendes *et al*., 2013) for details. (B) Heat map of kinematic parameters. Features are distinguished by “Step” (pink box), “Spatial” (yellow box), and “Gait” (black box), parameters. For each parameter values are matched to animals exposed to solvent (water) and *p* values are represented by a color code with red and blue shades indicate a decrease or increase relative to control, respectively. White indicates no variation. (C-F) PCA of all kinematic parameters. (C) Tridimensional representation of three-component PCA analysis. Each individual small dot represents one video while larger dots represents the average point (n=20 for each condition). Contribution of each component is indicated in each axis; (D) 2D representation with circles representing 50% of collected data; (E-F) Comparison of each PC coordinate. Boxplots represent the median as the middle line, with the lower and upper edges of the boxes representing the 25% and 75% quartiles, respectively; the whiskers represent the range of the full data set, excluding outliers (open circles). ***P < 0.001. (E) PC1 coordinates. (F) PC2 coordinates. See Methods for details.

### Time effect of BMAA-induced motor dysfunction

The reversibility of BMAA induced motor dysfunction or the effect of aging was subsequently verified. For this we evaluated kinematic performance using the FlyWalker system of 3-week old flies maintained on normal, untainted food, but exposed during their development to increasing concentrations of BMAA. Since *Drosophila* have a life expectancy of ∼60 days with a 50% survival rate of 40 days (Ashburner, 1989), we first tested if ageing *per se* could affect kinematic features. Interestingly, we found that 3-week-old flies display alterations in many parameters, including spatial parameters, consistent with age-related movement uncoordination (Fig. S5 and S6). Similarly to 1-week old animals, we found a dose-dependent effect of BMAA on kinematic features of this older age group (Fig. 5 and S7). Higher concentrations of BMAA induced a more significant uncoordinated stance profile (Fig. 5A) and larger differences when compared to control conditions (Fig. 5B). However, two noteworthy aspects should be highlighted. Firstly, compared to control animals, the impact of BMAA exposure in 3-week animals appears to be reduced when compared to 1-week animals. For example, exposure to 0.5 µM of BMAA during development affects many of the kinematic parameters in 1-week old animals (Fig. 4A), while in 3-week old animals only two parameters showed a significant difference to control conditions (Fig. 5 and S7). Secondly, while in 1-week old animals a seemingly linear dose dependence in response on kinematic effects can be observed (Fig. 4E and S4), this is not the case for 3-week-old animals (Fig. 5E and S7). These results suggest an adaptation of the neuromuscular system regarding the detrimental effect of BMAA during development. Nevertheless, at higher concentrations of BMAA (500 µM) this compensatory effect was less pronounced (Fig. 5B-E and S7), suggesting a threshold concentration for which the neurodevelopmental impairment no longer can be compensated and becomes permanent.

**Fig. 5.**
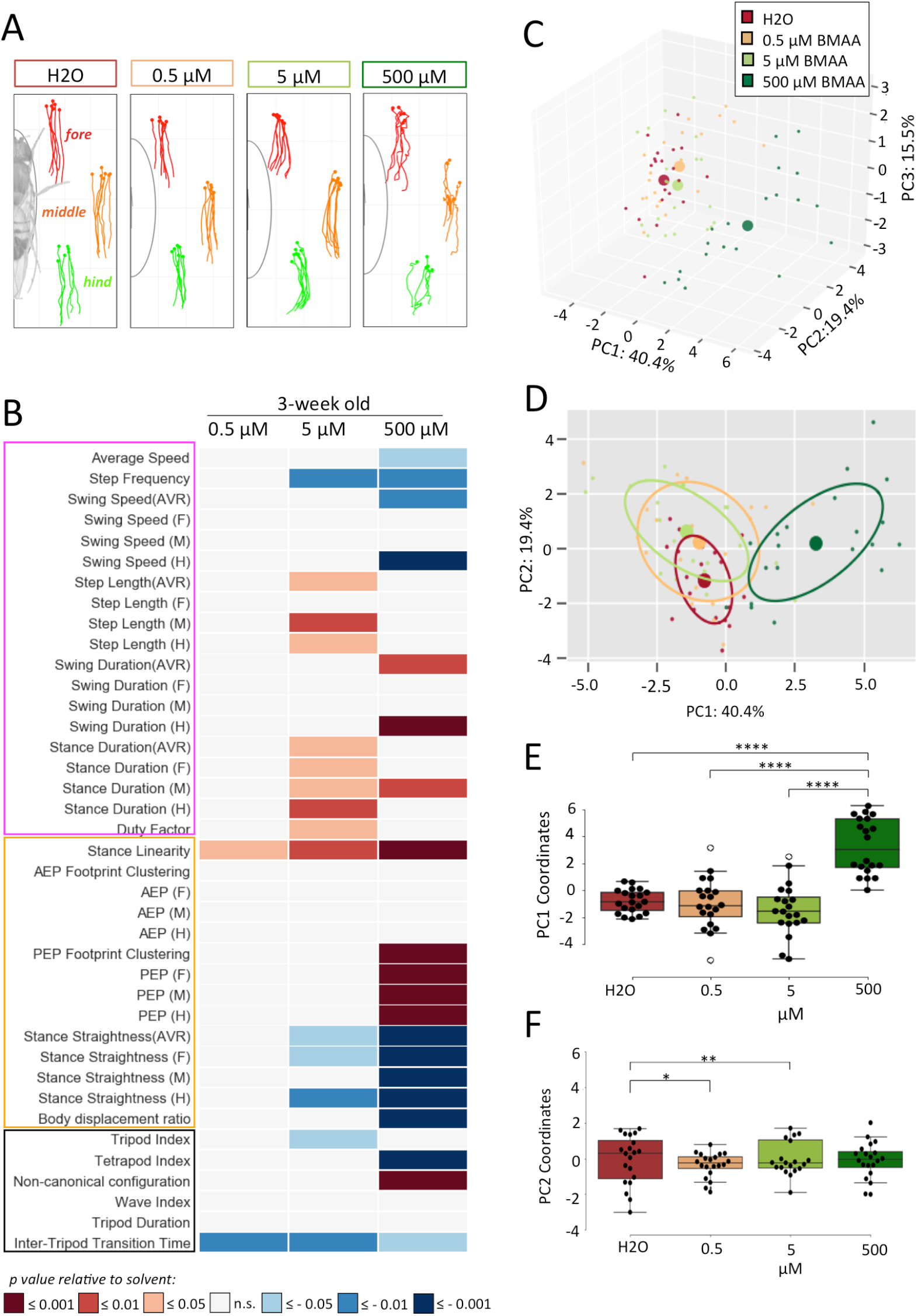
Kinematic parameters of 3-week adult animals exposed to increasing concentrations of BMAA during development. Exposure levels were 0; 0.5; 5 or 500 µM of BMAA. Kinematics features were recorded and analyzed using the FlyWalker system (n=20 for each condition). (A) Representative stance traces, which marks the tarsal contacts relative to the body axis during stance phases. See (Mendes *et al*., 2013) for details. (B) Heat map of kinematic parameters. Features are distinguished by “Step” (pink box), “Spatial” (yellow box), and “Gait” (black box), parameters. For each parameter values are matched to animals exposed to solvent (water) and *p* values are represented by a color code with red and blue shades indicate a decrease or increase relative to control, respectively. White indicates no variation. (C-F) PCA of all kinematic parameters. (C) Tridimensional representation of three-component PCA analysis. Each individual small dot represents one video while larger dots represents the average point (n=20 for each condition). Contribution of each component is indicated in each axis; (D) 2D representation with circles representing 50% of collected data; (E-F) Comparison of each PC coordinate. Boxplots represent the median as the middle line, with the lower and upper edges of the boxes representing the 25% and 75% quartiles, respectively; the whiskers represent the range of the full data set, excluding outliers (open circles). ***P < 0.001. (E) PC1 coordinates. (F) PC2 coordinates. See Methods for details.

**Fig. 7.**
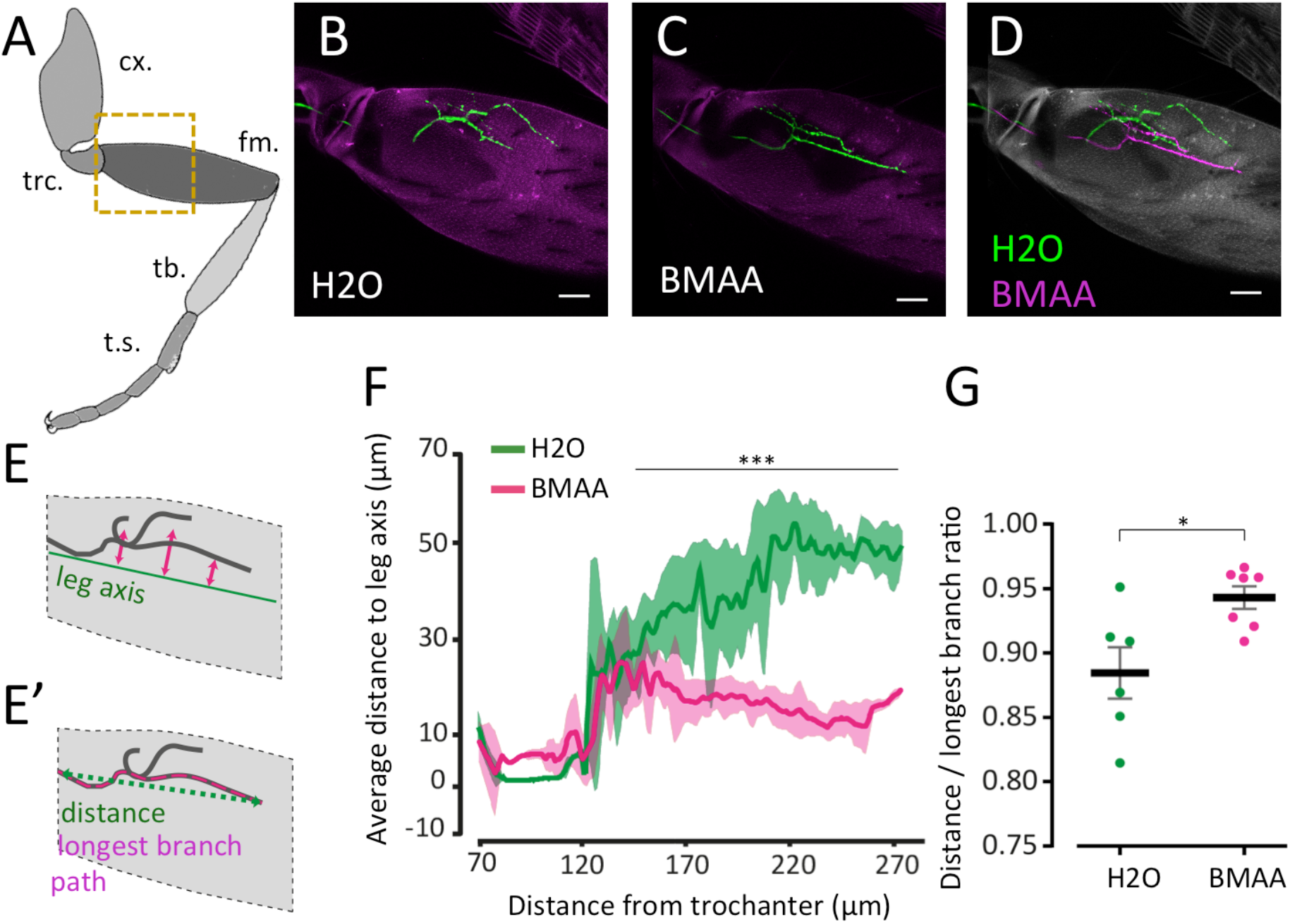
Long-term effect on motor neuron morphology by exposure to BMAA during development. Animals were exposed to solvent (water) and 500 µM of BMAA during development, and maintained in untainted food for three weeks before being examined. Genotype: ;R22A08-LexA/+; lexOp-Rab3:YFP/+. (A) Schematic representation of the fly leg. cx., coxa; trc., trochanter; fm., femur; tb., tibia; t.s., tarsal segments. Dashed square represents the scanned region in B-D. (B,C) Representative projections of a single motor neuron in the proximal femur (green) and cuticle autofluorescence (pink) in control animals (B) and exposed to 500 µM of BMAA during development (C). Bar, 30 µm. (D) Overlapped projections of the aligned images shown in B (green) and C (pink). Cuticle autofluorescence (white). Bar, 30 µm. (E) Schematic of the quantification presented in (F). The average distance to leg axis was calculated by measuring the distance between a line along the center of the femur to the average YFP signal perpendicular to the proximal-distal axis. (E’) Schematic of the quantification presented in (G). The distance to longest-branch ratio was measured by identifying the longest branch from the trochanter-femur joint and calculating the ratio between the distance of the longest branch and its straight line path. See methods for details. (F) Average distance to leg axis along the proximal-distal axis. Average values for animals developing in water (green, n=6) or BMAA (pink, n=7) are represented with shadowed areas representing standard deviation. Statistical analysis between 150 and 270 µm using Mann-Whitney non-parametric test, **** P < 0.0001). (G) Distance to longest branch ratio was calculated for animals developing in water (green, n=6) or BMAA (pink, n=7) Plots represent the mean as the middle line, with the error bars representing standard deviation. Statistical analysis was performed using Mann-Whitney non-parametric test, * P < 0.01.

Subsequently, we tested the hypothesis that the motor dysfunction induced by BMAA, particularly at higher concentrations, could include a time dependent neurodegenerative component beyond the initial neurodevelopmental effect. To test this hypothesis, we compared the behavioral 3-dimentional PCA of 1- and 3-week old animals exposed to increasing concentrations of BMAA during development (Fig. 6). Interestingly, we found that ageing did not exacerbate the differences between the control condition and flies reared in the presence of BMAA, even at the maximum concentration of 500 µM. These results support the model that BMAA interferes with the proper development of the neuromuscular system leading to an uncoordinated phenotype.

**Fig. 6.**
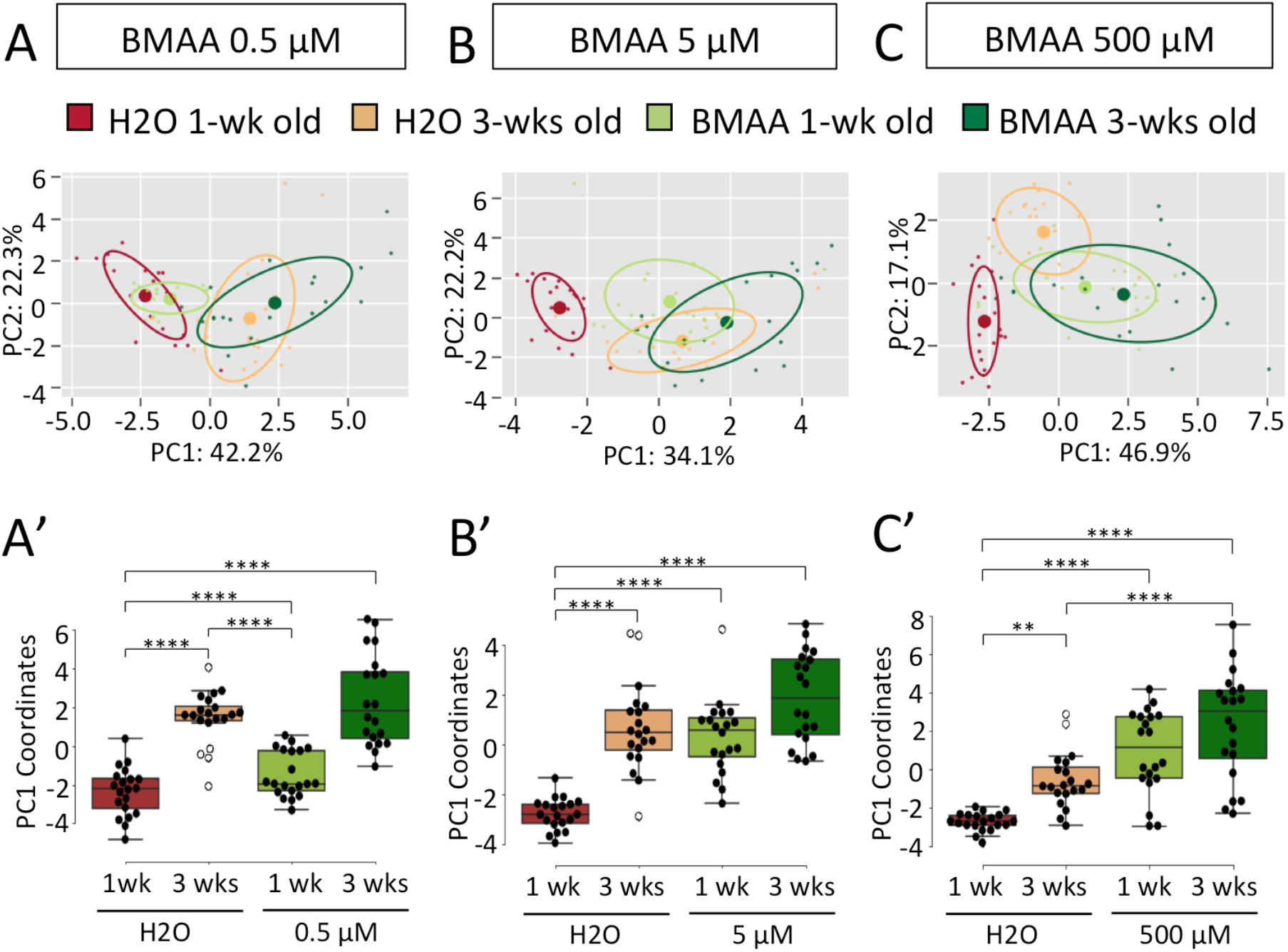
Time effect on motor dysfunction induced by BMAA neurotoxicity. (A-C) Animals were exposed to solvent (water); 0.5; 5; or 500 µM of BMAA during development, and maintained for one and three weeks before being tested for kinematic features using the FlyWalker system (n=20 for each condition). Data is represented as a 2D representation of a three-component PCA. Each individual small dot represents one video while larger dots represents the average point, with circles representing 50% of the data (n=20 for each condition). Contribution of each component is indicated in each axis (A’-C’) Comparison of PC1 coordinates. Boxplots represent the median as the middle line, with the lower and upper edges of the boxes representing the 25% and 75% quartiles, respectively; the whiskers represent the range of the full data set, excluding outliers (open circles). Statistical analysis with one-way ANOVA followed by Tukey’s *post hoc* test (for normal distributions) or Dunn’s *post hoc* test (for non-normal distribution), **P < 0.01, ***P < 0.001.

### Section 1.01 Effect of BMAA on motor neuron targeting

We found that exposure to BMAA during development induces motor impairments during adulthood (Figs 4-6 and S4-S7). Neurodevelopmental dysfunction of motor neurons has been shown to cause motor impairments with dendritic or axonal mistargeting, causing abnormal execution of motor commands (Enriquez *et al*., 2015). We hypothesized that the neuromuscular system is targeted by BMAA, resulting in an uncoordinated walking pattern. We thus tested if the motor defects induced by the developmental exposure to BMAA could be caused by a defect of the muscular fibers or a neurodevelopmental defect of the motor neurons.

We first tested if defects in muscular fibers were responsible for the motor phenotype observed. For this we used a *Drosophila* strain that expresses a GFP fusion-protein of Tropomyosin 1 (Buszczak et al., 2007), an integral component of actin filaments within muscle fibers (Lehman et al., 2009; Schachat et al., 1985; Wang et al., 1990), generating a muscle pattern that can be visualized by confocal microscopy (Fig. S8). Animals exposed to 500 µM BMAA did not demonstrate any defect in their tropomyosin1-GFP muscle pattern, both in general pattern and periodicity (Fig S8A-C), indicating that muscular dysfunction is not the cause of mobility coordination defects caused by BMAA.

We next questioned if development of motor neurons could be influenced by BMAA. To test this possibility we genetically labeled the neuromuscular junction (NMJ) of a single motor neuron enervating the tibia depressor muscle (tidm) within the proximal femur (Azevedo et al., 2020; Soler et al., 2004) (Fig. 7A). This was obtained by expressing the Rab3-YFP fusion protein that marks presynaptic sites (Zhang et al., 2007), driven by a motor neuron specific promoter (Azevedo *et al*., 2020). Three-week old control (solvent exposed) animals displayed a normal NMJ pattern, very similar to the one previously reported (Azevedo *et al*., 2020) (Fig. 7B). Strikingly, we found that exposure to 500 µM of BMAA during development induced an altered NMJ pattern, with projections located more dorsally, closer to the center of the femur, while targeting the same tidm muscle (Fig. 7C-F). Moreover, BMAA-exposed animals displayed a more linear axonal profile compared to control animals, which showed a more twisted appearance (Fig. 7E’,7G). It should be noted that other metrics, such as the number of branches and the most distal position, remain unchanged (Fig. S8D, E and data not shown). These results indicate that although animals developing in the presence of BMAA display an altered neuronal pattern, no signs of neurodegeneration were visible, consistent with the kinematic results observed previously (Fig. 6). As such, BMAA exposure during development can induce an altered motor neuron profile, which can ultimately be responsible for an impaired walking performance.

## Discussion

Here we present data on the use of a *Drosophila melanogaster* model for DNT studies, using a sensitive kinematic assay that accurately and quantitatively reports the status of the motor system. Our results indicate that this test strategy can identify changes in the neuro-muscular system that strongly matches the predicted neurotoxic effect of CPS, affecting several motor parameters without affecting survival during development (Fig. 1, 2, S2 and S3). Noteworthy, while we also observed a significant kinematic dysfunction in flies, exposed to DMSO-at general non-toxic concentrations during development (Fig. 2 and S3), we could resolve the kinematic effect of the additional presence of CPS further validating the sensitivity of our approach.

Moreover, we found that the putative neurotoxic agent BMAA induces strong motor defects in a dose dependent fashion in adult animals exposed to this compound during development, possibly by inducing motor neuron mistargeting, without inducing lethality (Fig. 3-7 and S4-S8). The sensitivity of the FlyWalker approach also identified kinematic degeneration and dysfunction in 3-week old, non-exposed flies, raising the possibility of using this system to identify age-related alterations in the motor system.

Exposure to neurotoxic agents can adversely interfere with the function and development of the nervous system in humans (Grandjean and Landrigan, 2006). Several chemical substances have been shown to induce DNT, given the susceptibility of the developing brain (Aschner *et al*., 2017). Nevertheless, neurotoxicity has also been reported in adult exposure, including environmental and in occupational settings. Known neurotoxic agents include mercury, lead, methylmercury, polychlorinated biphenyls, arsenic, and toluene (Grandjean and Landrigan, 2006; 2014). The assessment of neurotoxicity is usually carried out using OECD test guidelines and those of other national regulatory agencies (US EPA, guidelines), based on animal models. These models assess changes in neuroanatomical, neurophysiological, neurochemical and neurobehavioral parameters (Makris et al., 2009). However, there is a paucity of data on DNT. Of the 350,000 chemicals in use globally (Wang et al., 2020), DNT data is only available for approximately 110–140 compounds (Sachana et al., 2021). Current data requirements for *in vivo* DNT testing are considered insufficient to adequately screen and characterize compounds potentially hazardous for the human developing brain (Bal-Price et al., 2015a). Therefore, a pressing need exists for developing alternative methods that can more rapidly and cost-effectively support the identification and characterization of chemicals with DNT potential. Alternative DNT testing methods have been proposed, including human in vitro models, non-mammalian 3Rs models and *in silico* approaches (Bal-Price and Fritsche, 2018).

CPS, presents strong evidence for DNT effects in humans, and is listed as a reference compound for validation of alternative test methods to indicate DNT potential (Aschner *et al*., 2017). It is a chlorinated organophosphorus ester widely used as an insecticide in agricultural settings. CPS acts by inhibiting the enzyme acetylcholinesterase (AChE), thus preventing the degradation of the neurotransmitter acetylcholine (ACh) within synaptic clefts, both at neuromuscular or neuro-glandular junctions. This leads to ACh accumulation and cholinergic hyperstimulation (Casida, 2017). While CPS may elicit neurotoxic effects, CPS-oxon, resulting from cytochrome P450 mediated metabolism, is considered to be the primary neurotoxic agent. CPS is highly lipophilic and is readily absorbed through the skin via topical contact and lungs via inhalation. CPS is also quickly absorbed through the placenta and into fetal tissues, including the brain, during development. Exposure to CPS during pregnancy or early childhood has been implicated to cause learning and behavioural effects, including developmental delays related to cognition and motor function, attention deficit hyperactivity disorder, autism spectrum disorder (ASD), and tremors (Bai et al., 2014; Gray and Lawler, 2011; Rauh et al., 2011; Rauh et al., 2015; Rauh et al., 2012). CPS toxicity has been evaluated in several model systems, some replicating the neurotoxic effects observed in humans (Silva, 2020). Neuromotor function was impaired in rats and mice regardless of gestational, postnatal, or gestational plus postnatal exposure to CPS, while cognition was mostly decreased. Interestingly, qualitative results showed concordance among rodents, zebrafish and *C. elegans* for adverse effects on locomotor activity, neuromotor function, and AChE inhibition (Silva, 2020).

In our system, CPS displayed an impact on 16 motor parameters, most notably step and spatial parameters, suggesting that both motor and pre-motor centers at the level of the ventral nerve cord (the equivalent of the vertebrate spinal cord) are targeted leading to an uncoordinated walking behavior. Overall, the CPS data set demonstrates the high sensitivity and detailed motor information obtained with the FlyWalker approach on this known neurotoxic agent, largely coinciding with its known mode of action, with additional mechanistic clues.

While DMSO was used as a non-toxic solvent to prepare working solutions of CPS, the significant kinematic dysfunction observed was unexpected since there was no observed lethality at the concentration used. DMSO is an aprotic solvent that can solubilize a wide variety of otherwise poorly soluble polar and nonpolar molecules for which there are no alternative solvents (Santos et al., 2003); OECD Guidance Document on Good In Vitro Method Practices (GIVIMP). At commonly used concentrations, as was the case in our study, 70 mM corresponding to 0,5% (v/v), DMSO is not cytotoxic, and indeed has several pharmacological uses, such as an anti-inflammatory and reactive oxygen species scavenger (Santos *et al*., 2003), and is routinely used as a cryoprotectant in autologous bone marrow and organ transplantation. DMSO is considered a relatively safe solvent in doses up to 50 mg per day (Hanslick et al., 2009). Nevertheless, DMSO readily crosses the blood brain barrier, and has been reported to be neurotoxic. Hanslick et al. exposed C57Bl/6 mice of varying postnatal ages (P0–P30) to DMSO and observed widespread apoptosis in the developing mouse brain at all ages tested (Hanslick *et al*., 2009). The quantitative nature of the FlyWalker system allows to quantitatively compare different experimental conditions, delineating the simultaneous effect of two neurotoxic compounds, further validating our approach.

BMAA, is a natural non-proteinaceous amino acid produced by cyanobacteria (Cox et al., 2005), diatoms (Jiang et al., 2014) and dinoflagellates (Lage et al., 2014). Neurotoxicity of BMAA has been reported in various studies (Cox *et al*., 2005; Spencer et al., 1987). Particular interest in BMAA arose from the association with endemic neurodegenerative diseases, such as Parkinson-dementia complex and ALS, in the indigenous people of Guam (Kurland, 1988). Intense research efforts have identified various possible mechanisms of neurotoxicity, including misincorporation into cellular proteins, which may lead to adverse effects (Dunlop et al., 2013; Dunlop and Guillemin, 2019). BMAA is excitotoxic against neurons via glutamate receptors (Weiss and Choi, 1988), and also displays suppression of cell cycle progression of non-neuronal NIH3T3 cells via glutamate receptor-independent mechanisms (Okamoto et al., 2018).

None of the BMAA concentrations tested affected the survival rates (Fig. 3), eclosion timing or external morphology (not shown). However, increasing concentrations of BMAA rendered walking increasingly uncoordinated (Fig. 4 and S4), further suggesting a higher sensitivity of the developing neuronal system to toxic insults. This correlation between dosage and phenotype was attenuated in 3-week old animals, suggesting adaptation of the neuromuscular system to long-term motor constrains, leading to a shift in the walking behavior. Such adaptation has been described previously after long-term weight bearing or injury (Isakov et al., 2016; Mendes *et al*., 2014), underscoring a potential form of motor plasticity.

Our data indicates that BMAA exposure during development does not trigger acute adult neurodegeneration, instead promotes motor dysfunction phenotypically resembling aging. This conclusion is based on two observations: First, aged animals did not display enhanced effects of increasing BMAA concentrations in the uncoordination phenotype (3-weeks-old control animals resemble 1-week-old animals exposed to BMAA during development, see Fig. 6B’). Second, although the motor neuron analysis displayed an altered axonal pattern in animals raised on BMAA (Fig. 7), these do not show any metrics consistent with degeneration (Fig. 7 and S8).

It should be noted that our study focused on the effects of neurotoxic compounds during development, as adults were fed non-tainted food, and the reported neurodegenerative effects are probably due to a continuous exposure to BMAA (Kurland, 1988). Although it was out of the scope of this study, it would be interesting to test the long-term effects of BMAA in motor neurons degeneration and kinematic activity, or if alternatively, neurons become more susceptible to additional insults.

Finally, the observed effects of BMAA on motor neuron targeting suggests that the misincorporation of BMAA into proteins critical for the proper wiring of neuronal circuits such as transcriptional regulators or cell adhesion receptors can lead to altered terminal differentiation phenotypes ultimately leading to walking defects (Enriquez *et al*., 2015; Venkatasubramanian et al., 2019), and quite possibly other cognitive functions.

Besides CPS, the significant differences observed in the kinematic data obtained with DMSO and 3-week old flies highlight the sensitivity of the approach in measuring kinematic dysfunction. Exposure to neurotoxic agents may accelerate the dysfunction observed in aged individuals. Thus, overall, our results highlight the potential of detailed motor surveillance tools in simpler and more approachable model systems for the identification of neurotoxicity, with the potential to provide cues that can be explored in optimizing sensitive endpoints for pathway-specific screening and the determination of predictive key events.

## Author contributions

ASR, MK and CSM contributed to conception and design of the study. AC and CSM performed the experiments. AC, AMM and TP analyzed the raw data. AC, ASR, MK and CSM wrote the manuscript. All authors contributed to manuscript revision, read, and approved the submitted version.

## Acknowledgements

We thank Matthew Scott and Jun Zhang for the Rab3:YFP plasmid, John Tuthill for suggesting the R22A08 driver, Mendes lab, Rita Teodoro and her lab for comments during the execution of this project, Allan Mancoo for assistance in the design of the PCA script, and Daniela Pereira for comments on the manuscript. We also thank CONGENTO: consortium for genetically tractable organisms and the Bloomington Drosophila Stock Center for fly stocks. This work was supported by H2020 Marie Skłodowska-Curie Actions [H2020-MSCA-IF-2016, #752891, GEMiNI to CSM], Fundação para a Ciência e Tecnologia [IF/01154/2014/CP1252/CT0003 to CSM]. AMM was supported by a doctoral fellowship from FCT (PD/BD/128445/2017). SR and MK were partially supported by grant UID/BIM/0009/2020 for the Research Center for Toxicogenomics and Human Health-(ToxOmics) of the Fundação para a Ciência e Tecnologia, Portugal

## Declaration of interests

The authors declare no competing interests.

## Methods

### Fly strains

Wild type Canton S flies, [R22A08]-LexA, nSyb-GAL4 and Tm1[CC00578] (TropGFP) strains were obtained from the Bloomington stock center. The lexOp-Rab3:YFP strain was generated by excising the Rab3:YFP ORF (a gift from Matthew Scott) using NotI and XbaI restriction enzymes and cloning this fragment into a *LexOp* - MCS (pLOT) plasmid (Lai and Lee, 2006). Transgenic lines were generated by standard P-element-mediated transformation procedures in an *yw* background. Lines were selected based on strength and background expression. Flies were kept under constant temperature (25°C) and humidity (∼70%).

### Embryo collection

Ten females and five males of wild type CantonS flies were transferred to a cage with a petri dish containing a layer of apple juice and maintained overnight. Produced eggs were collected and transferred to a 70 μm nylon cell strainer (BD Biosciences) with the help of distilled water and a brush. Embryos were submerged in a 50% (v/v) solution of commercial bleach for a few minutes, in order to dechorionate the eggs to test for fertilization. To identify fertilization and viability, each egg was individually observed with a fluorescence stereoscope (SteREO V8, Zeiss) with an Intermediate LED tube FL S, 38 HE GFP - (EX BP 470/40, BS FT 495, EM BP 525/50). Fertilized eggs were selected based on the presence of developing intestine and movement, distinguishable from non-fertilized ones, which showed a homogeneous white colouring. The former ones were transferred from the container for further experimentation.

### Contaminated food preparation

Eggs were exposed to contaminated and control food in vials containing 1 gram of food (approximately 1 ml) in a 2 cm diameter vial. For each condition, stock solutions of test compounds were prepared and 5 µL added to a vial of food, except in the case of DMSO (see below). Food was mixed using a handheld drill with a disposable fork with two tines. To obtain a homogenous layer, the vials were centrifuged at 280 g. Lastly, food was left overnight to allow even spreading of test compound by diffusion. For each condition, 3 vials were prepared.

Control experiments were carried out to evaluate potential neurotoxic effects of the solvent DMSO. Three food concentrations were studied, namely 0.07, 70 and 140 mM. The two highest concentrations were obtained by adding 5 and 10 µL, respectively, of pure DMSO (Merk, Darmstadt, Germany) into the food vial. For 0.07 mM, pure DMSO was diluted 100× in water and 5 µL were added to the food vial.

Regarding CPS (Sigma-Aldrich, St. Louis, MO, USA), a stock solution of 1 M was prepared in DMSO. This solution was further diluted in DMSO to obtain 10, 1 and 0.1 mM, and 5 µL of these solutions were added to fly food vials to generate 50, 5 and 0.5 µM, respectively, leading to a constant DMSO of 70 mM (0.5% v/v) in each of these three dose levels.

The amino acid BMAA (Sigma-Aldrich, St. Louis, MO, USA), was dissolved using water as solvent to obtain a stock solution of 1 M. This solution was further diluted to obtain 1000, 100, 10, 1 and 0.1 mM, and 5 µL of these stock solutions were added to fly food vials to obtain 5000, 500, 50, 5 and 0.5 µM final food concentrations, respectively.

Adult animals were kept on normal, untainted food for the FlyWalker assay and imaging experiments. If necessary, food was changed every week.

### Survival and pupal climbing positioning

For each exposure condition, at least 25 fertilized and viable eggs were placed in each test vial, performed in triplicate, totalling at least 75 animals. The positioning of the pupae in the vial wall were measured according to four height-locations: I. pupae located inside the food; II. placed slightly above the food; III. positioned up to 1 centimetre (cm) from the food; and IV. occupying the area above 1 cm (Fig. 1D). The number of animals reaching pupal stages and eclosing (exiting their pupal cases) were additionally recorded.

### Kinematic analysis

Kinematic behavioural experiments were carried out as described previously (Mendes *et al*., 2013). After eclosion, flies were collected and kept for 3 days in a new vial containing normal fly food and a humid paper filter, in order to prevent contamination of the walking arena. Individual flies were placed into a walking chamber and filmed with a Photron (Tokyo, Japan) Mini UX-100 camera using a Nikon (Tokyo, Japan) AF 24-85mm lens at 250 frames per second (fps). For each condition, 10 animals were filmed twice, generating 20 videos. Kinetic parameters of fly movement were obtained through analyses of obtained videos, using the FlyWalker software package (Mendes *et al*., 2013).

In addition to the previously published kinematic parameters (Mendes *et al*., 2013; Mendes *et al*., 2014), two additional parameters were quantified:

### Stance Straightness

Ratio between the distance from AEP to PEP (stance phase) and the path described by the tarsal contacts relative to the body (stance trace).

### Body displacement ratio

Ratio between of the distance traveled over the body path.

### Leg dissection and microscopy

T1 legs were isolated using a dissecting microscope from selected animals with the correct genotype. Legs were kept in cold PBS, followed by a quick rinse with ethanol to remove the wax from the cuticle, preventing floating and maintaining legs submerged during the procedure. After three washes in PBS, legs were fixed overnight in 4% paraformaldehyde in PBS at 4°C. After three washes in PBS, legs were mounted on glass slides in glycerol and imaged with a Zeiss (Oberkochen, Germany) LSM710 confocal microscope with a 40× objective using a 514 nm excitation laser and a emission windows of 519-582 nm (for YFP), and a 594 nm excitation laser and a emission windows of 599-797nm (for auto-fluorescence).

TropGFP images were analysed using Fiji for profile tracing (Schindelin et al., 2012), and a MATLAB (MathWorks Inc, Natick, MA, EUA) custom script for peak signal measurement and coefficient of variation calculation. Muscle profiles were obtained using a Fiji line tool manually perpendicular to each muscle fibre in locations with good signal in different locations of 4 independent images of for each condition. Muscle profiles were then analysed using a custom MATLAB script, which detected each peak (“find-peaks” function) and quantified the periodicity of each profile. The coefficient of variation (standardized measure of dispersion of a probability distribution) of each profile was then calculated and plotted.

Motor neuron imaging data was analysed using a custom MATLAB script. Leg upper and lower boundaries were segmented using the cuticle autofluorescence signal. Noise was reduced using a gaussian filter function (imgaussfilt) with sigma 2 and a global threshold was determined. With these two boundaries a central leg axis was detected and used for subsequent calculation. The neuronal profile was binarized using an OTSU global threshold and the intensity centre of mass in each position was found along the proximal-distal axis.

For the ratio between NMJ penetration depth (starting from trochanter-femur joint) and the total length of the longest branch (starting from trochanter-femur joint) images were processed in Imaris 9.5 (Oxford instruments, Abingdon, U.K.) using the filament tracer tool semi-automatically (results were manually curated) and measurements were made using the measurement and statistics tools.

All other measurements were carried out using the Imaris 9.5 imaging analysis package.

### Data analysis

Kinematic parameters were extracted using the Flywalker Software, in which each video resulted a single data point (Mendes *et al*., 2013). Since many of the measured gait parameters vary with speed, we analysed the data for these parameters by firstly determining the best-fit regression model for the control experiment. Subsequently the residual values for each experimental group in relation to this regression model were determined, using RStudio 1.1.442 (Mendes *et al*., 2014). The residual data set was subsequeently tested for normality and homoscedasticity using the Shapiro-Wilk and Levene Tests. Statistical significance between the control and each of the experimental groups were determined using Kruskal-Wallis analysis of variance followed by Dunn’s post hoc test (for non-normal distributions) or one-way-Anova followed by Tukey’s post hoc test (for normal distributions). Significant differences were represented by a Heatmap where each column represents the different groups compared with the control and in each line the kinematic parameters. Red and blue bars indicate an increased and decreased value, respectively. Lighter, intermediate, and darker colours indicate a p-value of < 0.05, < 0.01 and < 0.001, respectively. Boxplots represent the median as the middle line, with the lower and upper edges of the boxes representing the 25% and 75% quartiles, respectively. The whiskers represent the range of the full data set, excluding outliers. Outliers are defined as any value that is 1.5 times the interquartile range below or above the 25% and 75% quartiles, respectively. Statistical analysis was performed using custom Python scripts and GraphPad Prism.

To have a more succinct representation of the data, facilitating its visualization, an unsupervised dimensionality reduction method was used, namely Principal Component Analysis (PCA). The data was first pre-processed by centering and scaling after which the PCA algorithm was applied. We chose the first three principal components to visualize the data. The first two components were chosen to generate a two-dimensional plot. Each dot in the plots corresponds to a video, representing these new abstract variables. Color-coded dots were used to distinguish specific groups. As such clusters of dots reflects similar walking patterns, shared by the corresponding flies.

## Supplementary figures

**Fig. S1.**
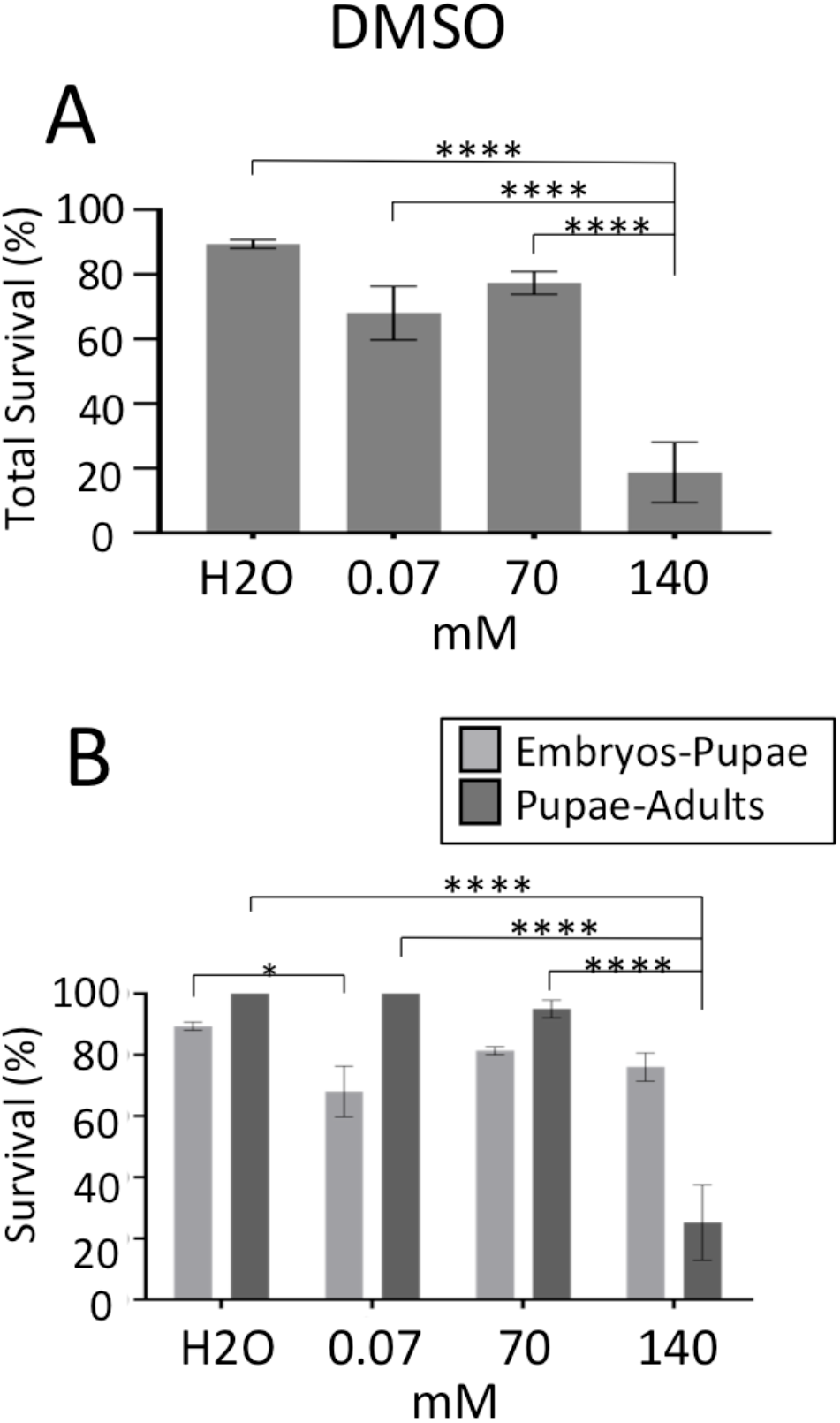
Pupal and adult survival of *Drosophila melanogaster* developed from embryos exposed to food tainted with DMSO. (A) Survival of adult animals after exposure to increasing concentrations of DMSO (0; 0.07; 70; or 140 mM, n=75 for each condition) compared to solvent (water). (B) Partial survival scores. Light grey represent embryo to pupae survival, and dark grey represent pupae to adult survival. Bar graphs represent the average percentage of animals surviving each developmental stage ± SEM. Statistical analysis with one-way ANOVA followed by Tukey’s *post hoc* test (for normal distributions) or Dunn’s *post hoc* test (for non-normal distribution), *P < 0.05; **P < 0.01; ***P < 0.001.

**Fig. S2.**
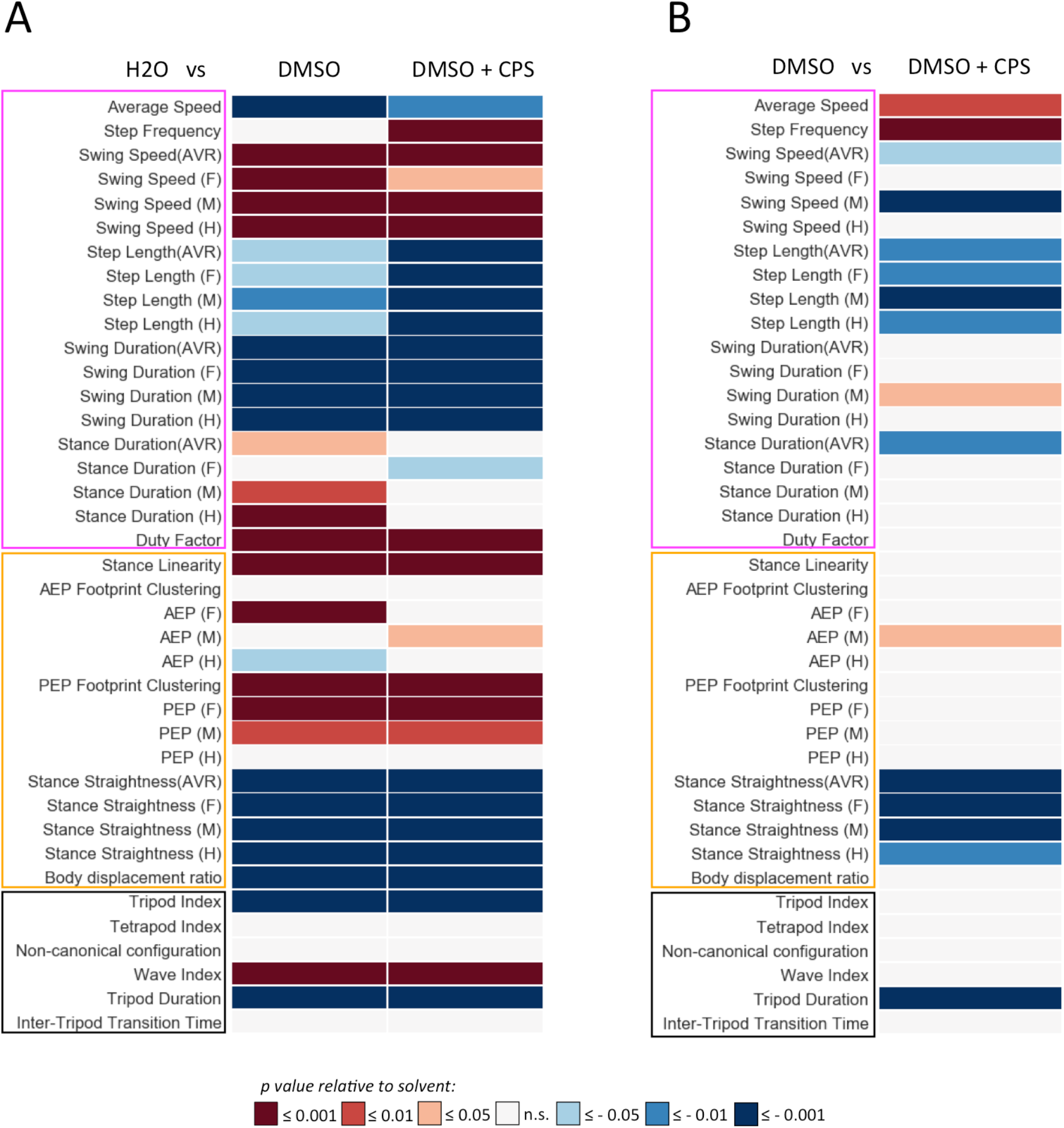
Heat map of kinematic parameters of adult animals exposed during development to water, 70 mM DMSO (solvent) and DMSO plus 5 µM CPS. Features are distinguished by “Step” (pink box), “Spatial” (yellow box), and “Gait” (black box), parameters. For each parameter values are matched to animals exposed to (A) water or (B) DMSO. *p* values are represented by a color code with red and blue shades indicate a decrease or increase relative to control, respectively. White indicates no variation.

**Fig. S3.**
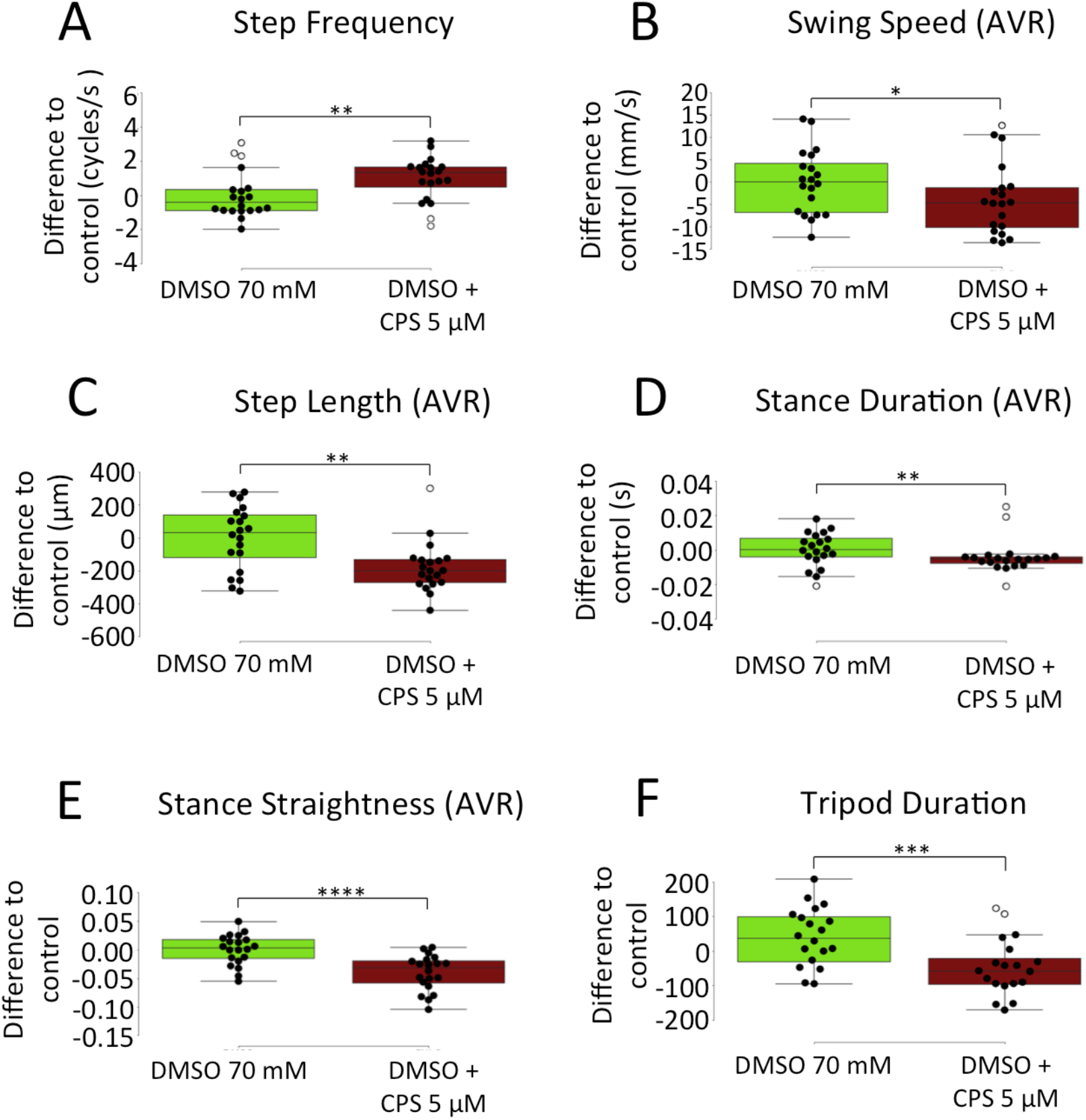
Representative kinematic parameters affected by exposure to 5 µM of CPS during development. Boxplots represent the median as the middle line, with the lower and upper edges of the boxes representing the 25% and 75% quartiles, respectively; the whiskers represent the range of the full data set, excluding outliers (open circles). Statistical analysis with one-way ANOVA followed by Tukey’s *post hoc* test (for normal distributions) or Dunn’s *post hoc* test (for non-normal distribution), *P < 0.05; **P < 0.01; ***P < 0.001. (A) Step frequency. (B) Swing speed. (C) Step length. (D) Stance duration. (E) Stance straightness. (F) Tripod duration.

**Fig. S4.**
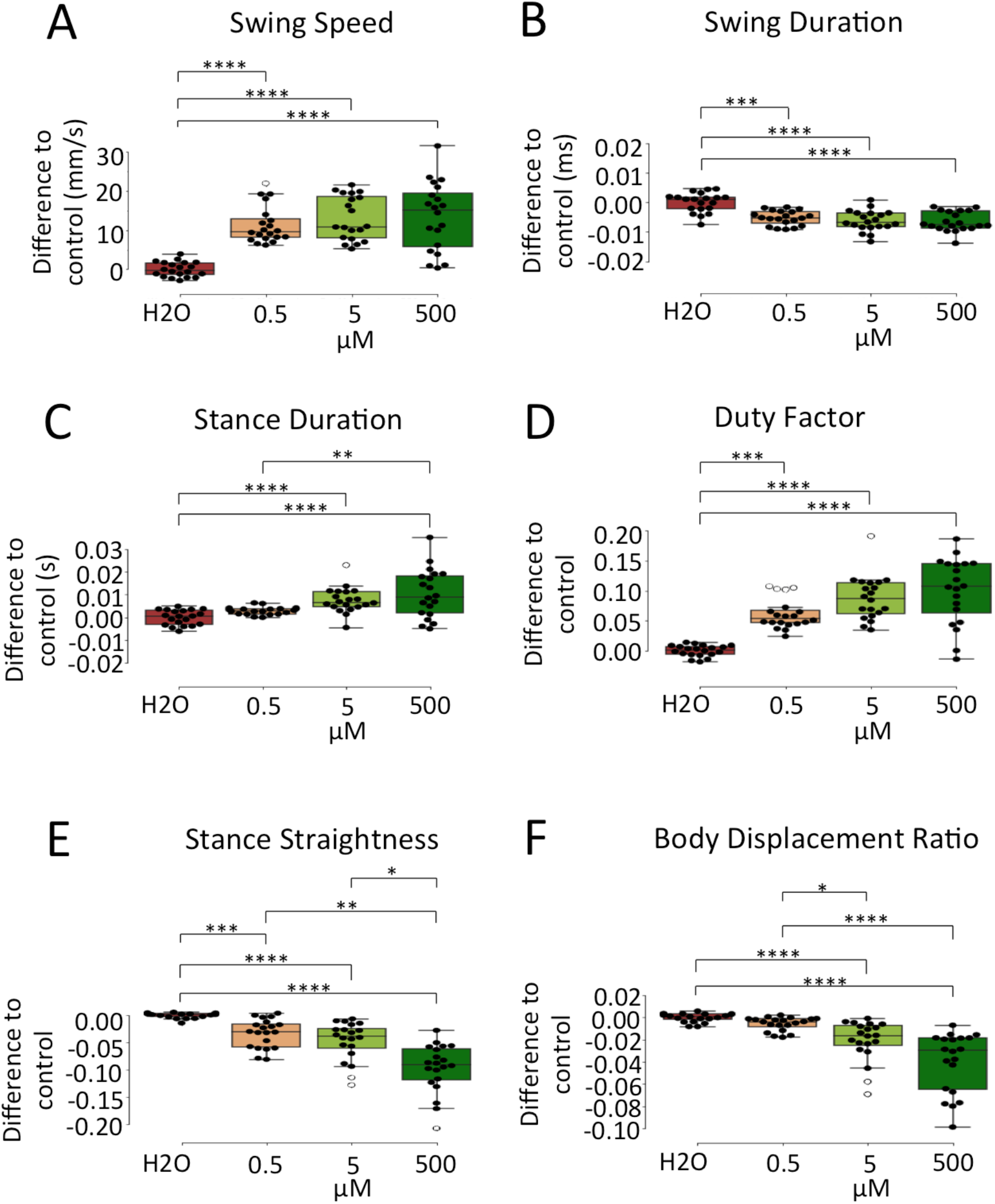
Representative kinematic effects of 1-week old animals exposed to increasing concentrations of BMAA during development. BMAA exposure levels were 0; 0.5; 5 or 500 µM. Kinematic features were recorded and analyzed using the FlyWalker system (n=20 for each condition). Boxplots represent the median as the middle line, with the lower and upper edges of the boxes representing the 25% and 75% quartiles, respectively; the whiskers represent the range of the full data set, excluding outliers (open circles). Statistical analysis with one-way ANOVA followed by Tukey’s *post hoc* test (for normal distributions) or Dunn’s *post hoc* test (for non-normal distribution), *P < 0.05; **P < 0.01; ***P < 0.001. (A) Swing speed. (B) Swing duration. (C) Stance duration. (D) Duty factor. (E) Stance straightness. (F) Body displacement ratio.

**Fig. S5.**
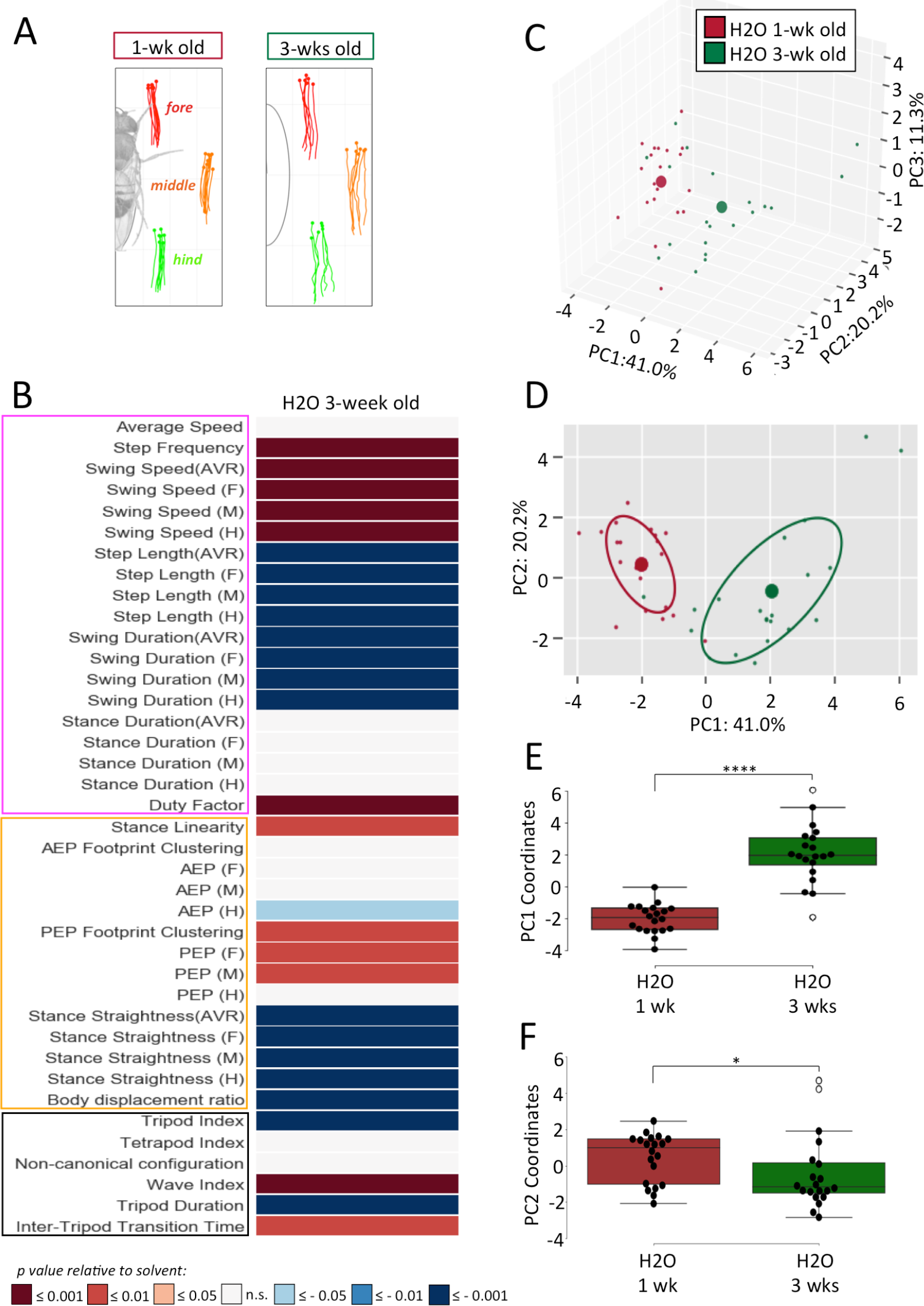
Kinematic effects of 1 vs 3-week old animals. (A) Representative stance traces of 1 and 3-week old animals grown in untainted food. (B) Heat map of kinematic parameters of 1 vs 3-week old wild type animals. Animals were kept on untainted food for 1 and 3 weeks and kinematic features were recorded and analyzed using the FlyWalker system (n=20 for each condition). Features are distinguished by “Step” (pink box), “Spatial” (yellow box), and “Gait” (black box), parameters. For each parameter values are matched to 1-week old animals and *p* values are represented by a color code with red and blue shades indicate a decrease or increase relative to control, respectively. Lighter, intermediate and darker shades indicate a p value ≤ 0.05; 0.01; and 0.001, respectively. White indicates no variation. See Methods for details. (C-F) PCA of all kinematic parameters of 1 and 3-week old animals. (C) 3D-representation of three-component PCA analysis. Each individual small dot represents one video while larger dots represents the average point (n=20 for each condition). Contribution of each component is indicated in each axis (D) 2D representation with circles representing 50% of collected data. (E) PC1 coordinates. (F) PC2 coordinates. Boxplots represent the median as the middle line, with the lower and upper edges of the boxes representing the 25% and 75% quartiles, respectively; the whiskers represent the range of the full data set, excluding outliers (open circles). Statistical analysis with one-way ANOVA followed by Tukey’s *post hoc* test (for normal distributions) or Dunn’s *post hoc* test (for non-normal distribution). *P < 0.05; ***P < 0.001.

**Fig. S6.**
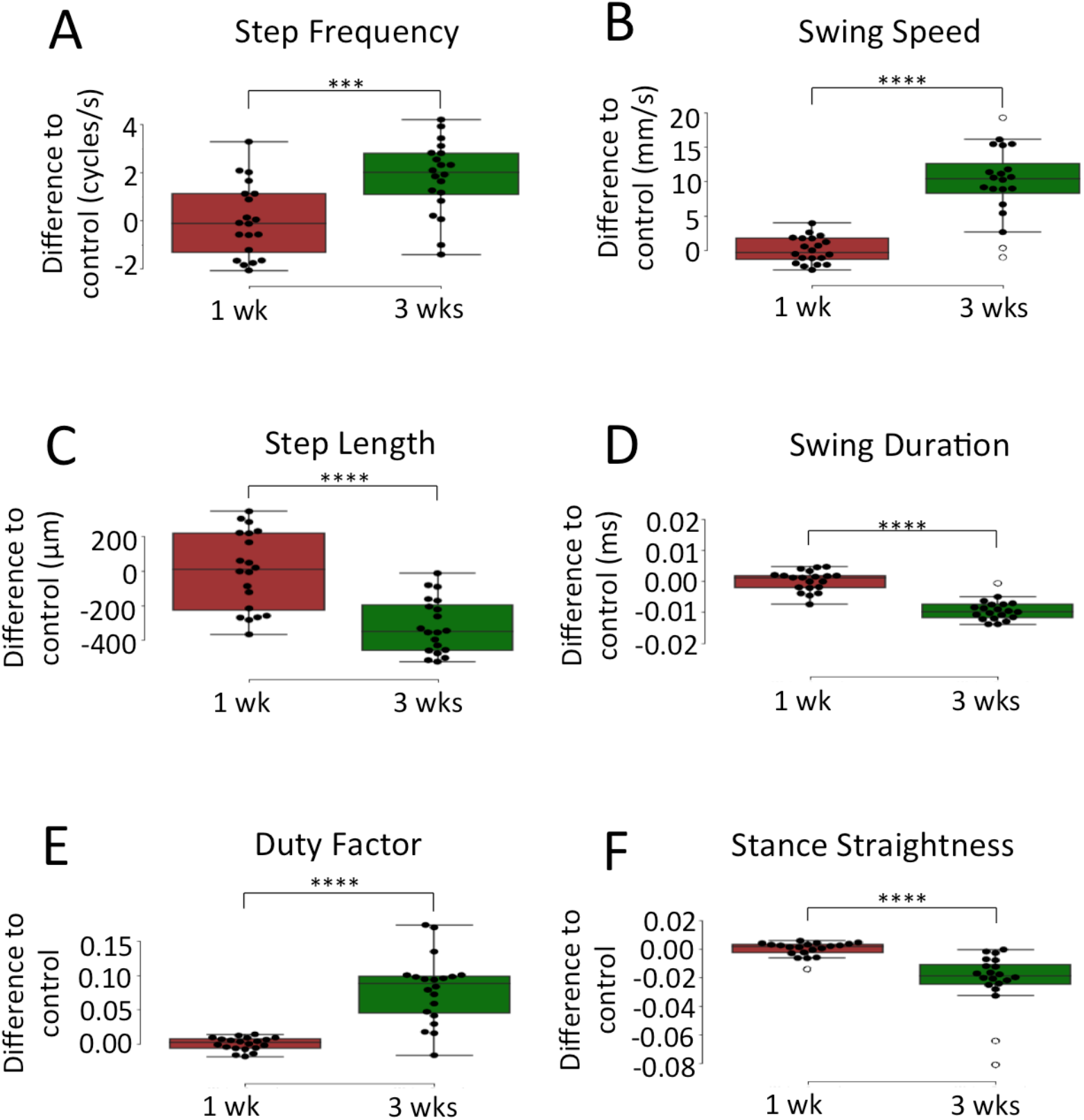
Representative kinematic parameters comparing 1 to 3-week old animals. Boxplots represent the median as the middle line, with the lower and upper edges of the boxes representing the 25% and 75% quartiles, respectively; the whiskers represent the range of the full data set, excluding outliers (open circles). Statistical analysis with one-way ANOVA followed by Tukey’s *post hoc* test (for normal distributions) or Dunn’s *post hoc* test (for non-normal distribution), ***P < 0.001. (A) Step frequency. (B) Swing speed. (C) Step length. (D) Swing duration. (E) Duty factor. (F) Stance straightness.

**Fig. S7.**
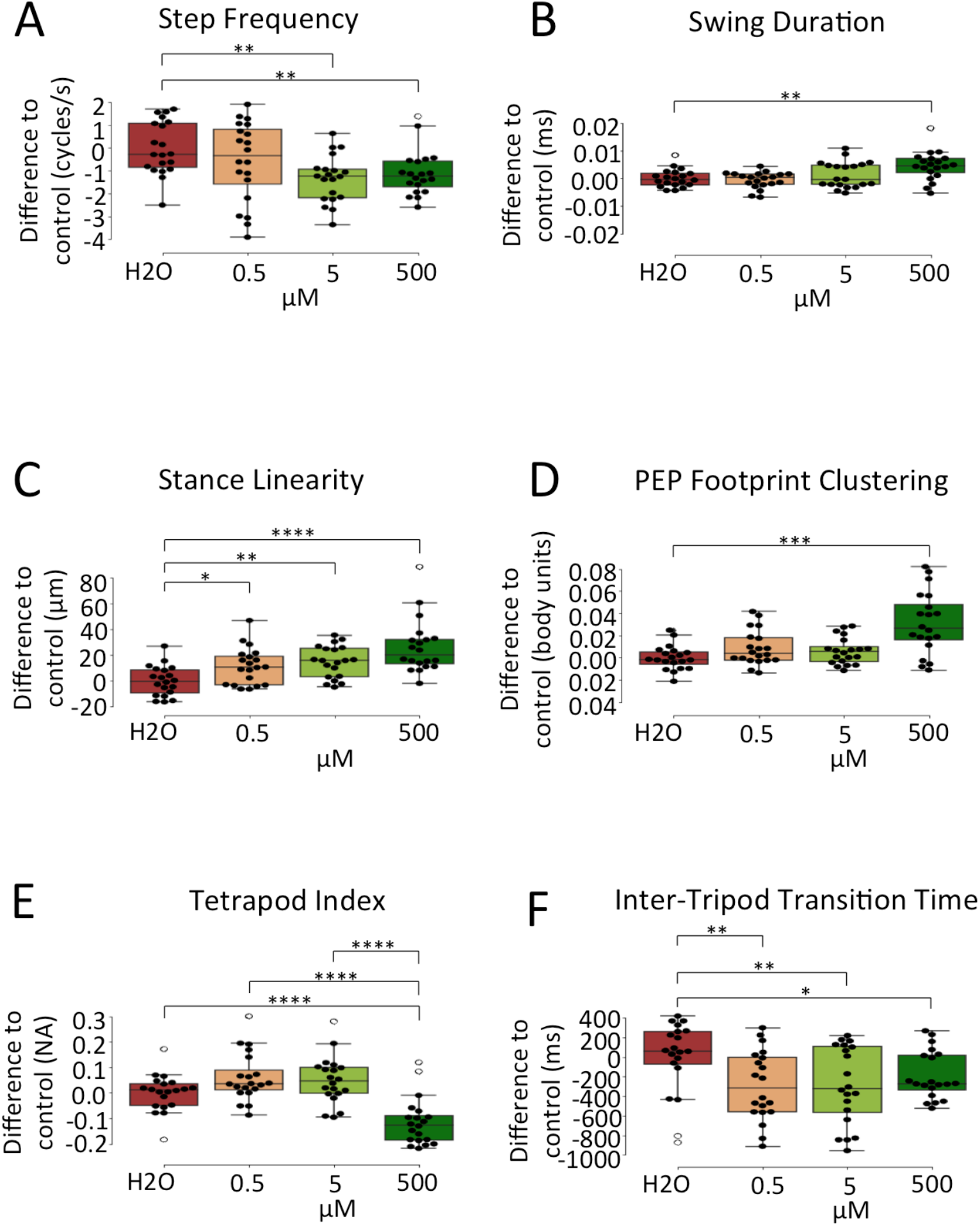
Representative kinematic effects of 3-week old animals exposed to increasing concentrations of BMAA during development. BMAA exposure levels were 0; 0.5; 5 or 500 µM. Kinematic features were recorded and analyzed using the FlyWalker system (n=20 for each condition). Boxplots represent the median as the middle line, with the lower and upper edges of the boxes representing the 25% and 75% quartiles, respectively; the whiskers represent the range of the full data set, excluding outliers (open circles). Statistical analysis with one-way ANOVA followed by Tukey’s *post hoc* test (for normal distributions) or Dunn’s *post hoc* test (for non-normal distribution), *P < 0.05; **P < 0.01; ***P < 0.001. (A) Step frequency. (B) Swing duration. (C) Stance linearity. (D) PEP footprint clustering. (E) Tetrapod index. (F) Inter-tripod transition time.

**Fig. S8.**
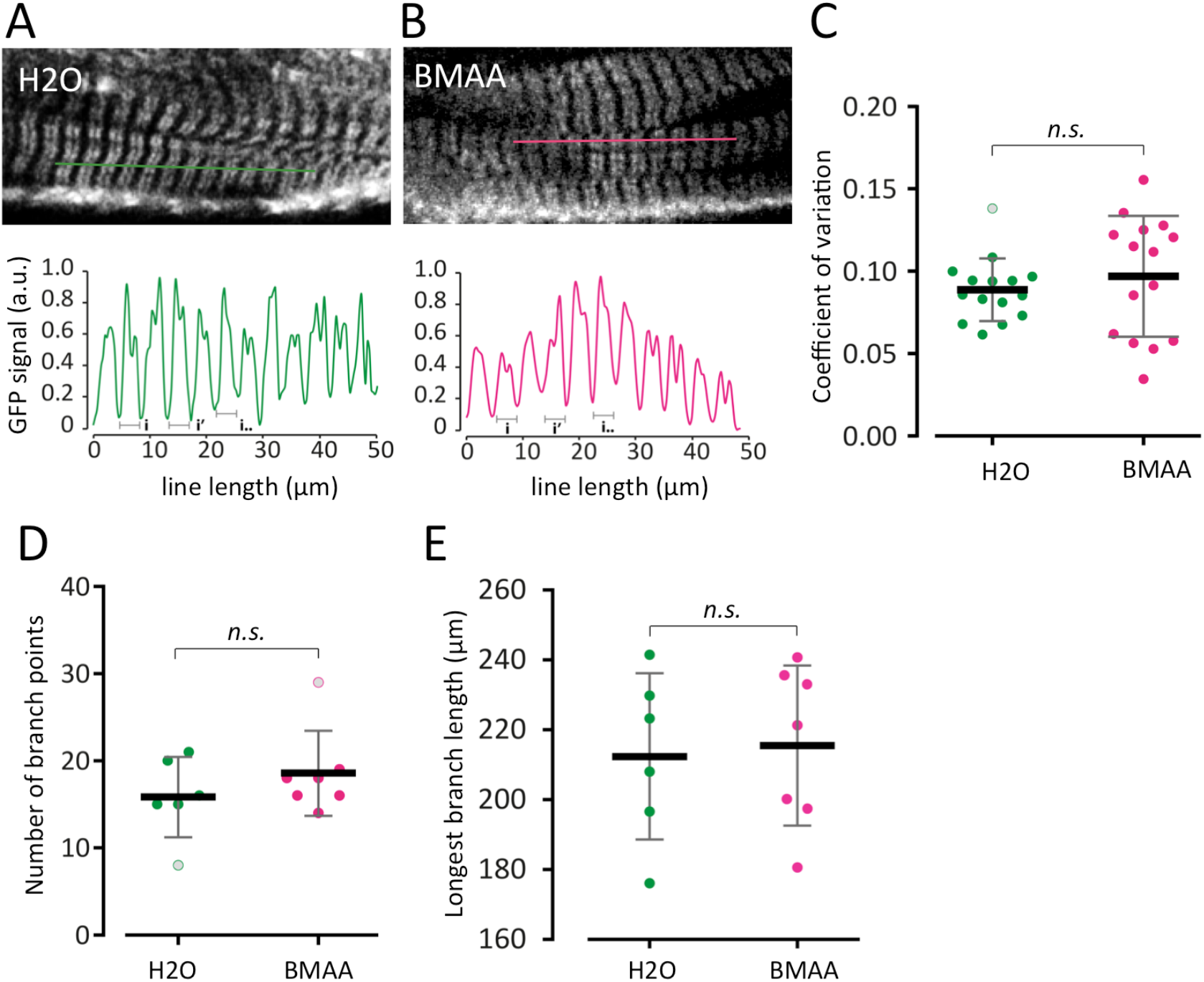
Long-term effect on muscle and motor neuron morphology by exposure to BMAA during development. Animals were exposed to solvent (water) and 500 µM of BMAA during development, and maintained in untainted food for three weeks before being examined. (A-C) Tropomyosin-GFP pattern in leg musculature. Genotype: Tm1[CC00578]/+ . (A-B) Upper panels show representative confocal sections of leg muscle patterns of animals exposed to (A) solvent (water) or (B) 500 µM BMAA (n=15 for each condition). Lines represent the region used to determine the signal intensity perpendicular to the muscle fiber. Lower panels show the variation of signal across the muscle fiber with lower inflexion points used to determine the periodicity of each profile (i). See Methods for details. (C) Coefficient of variation (standardized measure of dispersion of a probability distribution) between animals exposed to water and BMAA. (D,E) Quantification of the number of branch points (D), and longest branch length (E) in animals exposed to water (n=6) and 500 µM of BMAA (n=7) during development and maintained in untainted food for three weeks expressing Rab3:YFP under the control of the R22A08 driver. Genotype: ;R22A08-LexA/+; lexOp-Rab3:YFP/+. Plots represent the mean as the middle line, with the error bars representing standard deviation. Statistical analysis was performed using Mann-Whitney non-parametric test, n.s., non-significant.

**Table S1.**
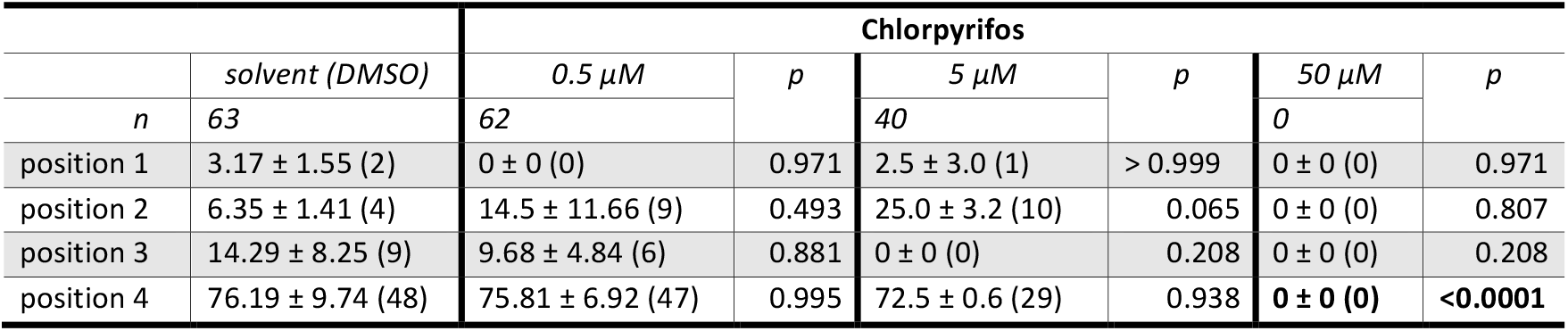
Pupal positioning of animals developing in the presence of solvent (DMSO); 0.5 µM; 5 µM; or 50 µM of chlorpyrifos. Values are expressed as a percentage of pupae in each position ±SEM. Number of animals analysed are indicated (n) and the number of pupas assigned to each position is indicated between parenthesis. Statistical analysis with one-way ANOVA followed by Tukey’s post hoc test (for normal distributions) or Dunn’s post hoc test (for non-normal distribution). Conditions that display statistically significant differences to control conditions are highlighted in bold. See Methods and main text for details.

**Table S2.** Pupal positioning of animals developing in the presence of solvent (water); 0.5 µM; 5 µM; 50 µM; 500 µM or 5000 µM of BMAA. Values are expressed as a percentage of pupae in each position ±SEM. Number of animals analysed are indicated (n) and the number of pupas assigned to each position is indicated between parenthesis. Statistical analysis with one-way ANOVA followed by Tukey’s post hoc test (for normal distributions) or Dunn’s post hoc test (for non-normal distribution). Conditions that display statistically significant differences to control conditions are highlighted in bold. See Methods and main text for details.

|  |  | BMAA |  |  |  |  |  |  |  |  |  |
| --- | --- | --- | --- | --- | --- | --- | --- | --- | --- | --- | --- |
| | solvent<br>(water) | 0.5 μM | p | 5 μM | p | 50 μM | p | 500 μM | p | 5000 μM | p |
| n | 67 | 60 |  | 55 |  | 72 |  | 65 |  | 63 |  |
| position 1 | 6.0 ± 1.6 (4) | 0 ± 0 (0) | 0.889 | 0 ± 0 (0) | 0.889 | 1.4 ± 1.4 (1) | 0.9628 | 0 ± 0 (0) | 0.889 | 0 ± 0 (0) | 0.889 |
| position 2- | 23.9 ± 7.4 (16) | **5.0 ± 2.8 (3)** | **0.011** | **5.4 ± 3.4 (3)** | **0.019** | **0 ± 0 (0)** | **0.001** | **4.6 ± 2.6 (3)** | **0.011** | 11.1 ± 2.8 (7) | 0.184 |
| position 3- | 13.4 ± 7.0 (9) | 36.7 ± 6.9 (22) | **0.001** | 36.4 ± 5.0 (20) | **0.002** | 13.9 ± 3.4 (10) | >0.999 | 4.6 ± 0.2 (3) | 0.573 | 20.6 ± 3.0 (13) | 0.815 |
| position 4- | 56.7 ± 3.4 (38) | 58.3 ± 4.3 (35) | 0.999 | 58.2 ± 7.0 (32) | 0.999 | **84.7 ± 2.4 (61)** | **<0.0001** | **90.8 ± 2.5 (59)** | **<0.0001** | 68.3 ± 5.3 (43) | 0.239 |

## Notes

### Competing Interest Statement

The authors have declared no competing interest.

